# Genome-wide association study of asthma in individuals of African ancestry reveals novel asthma susceptibility loci

**DOI:** 10.1101/112953

**Authors:** Michelle Daya, Nicholas Rafaels, Sameer Chavan, Henry Richard Johnston, Aniket Shetty, Christopher R. Gignoux, Meher Preethi Boorgula, Monica Campbell, Pissamai Maul, Trevor Maul, Candelaria Vergara, Albert M. Levin, Genevieve Wojcik, Dara G. Torgerson, Victor E. Ortega, Ayo Doumatey, Maria Ilma Araujo, Pedro C. Avila, Eugene Bleecker, Carlos Bustamante, Luis Caraballo, Georgia M. Dunston, Mezbah U. Faruque, Trevor S. Ferguson, Camila Figueiredo, Jean G. Ford, Pierre-Antoine Gourraud, Nadia N. Hansel, Ryan D. Hernandez, Edwin Francisco Herrera-Paz, Eimear E. Kenny, Jennifer Knight-Madden, Rajesh Kumar, Lesli A. Lange, Ethan M. Lange, Antoine Lizee, Alvaro Mayorga, Deborah Meyers, Dan L. Nicolae, Timothy D. O’Connor, Ricardo Riccio Oliveira, Christopher O. Olopade, Olufunmilayo Olopade, Zhaohui S. Qin, Charles Rotimi, Harold Watson, Rainford J. Wilks, L. Keoki Williams, James G. Wilson, Carole Ober, Esteban G. Burchard, Terri H. Beaty, Margaret A. Taub, Ingo Ruczinski, Rasika Ann Mathias, Kathleen C. Barnes, Ayola Akim Adegnika, Ganiyu Arinola, Ulysse Ateba-Ngoa, Gerardo Ayestas, Adolfo Correa, Francisco M. De La Vega, Celeste Eng, Said Omar Leiva Erazo, Marilyn G. Foreman, Cassandra Foster, Li Gao, Jingjing Gao, Kimberly Gietzen, Leslie Grammer, Linda Gutierrez, Mark Hansen, Tina Hartert, Yijuan Hu, Kwang-Youn A. Kim, Pamela Landaverde-Torres, Javier Marrugo, Beatriz Martinez, Rosella Martinez, Luis F. Mayorga, Delmy-Aracely Mejia-Mejia, Catherine Meza, Solomon Musani, Shaila Musharoff, Oluwafemi Oluwole, Maria Pino-Yanes, Hector Ramos, Allan Saenz, Steven Salzberg, Maureen Samms-Vaughan, Robert Schleimer, Alan F. Scott, Suyash S. Shringarpure, Wei Song, Zachary A. Szpiech, Raul Torres, Gloria Varela, Olga Marina Vasquez, Lorraine B. Ware, Maria Yazdanbakhsh on behalf of CAAPA

**Affiliations:** Department of Medicine, University of Colorado Denver, Aurora, CO, 80045, USA; Department of Biostatistics and Bioinformatics, Emory University, Atlanta, GA 30322, USA; Department of Genetics, Stanford University School of Medicine, Stanford, CA 94305, USA; Genetics and Epidemiology of Asthma in Barbados, The University of the West Indies, Chronic Disease Research Centre, Jemmots Lane, St. Michael, BB11115, Barbados; Department of Medicine, Johns Hopkins University, Baltimore, MD 21224, USA; Department of Public Health Sciences, Henry Ford Health System, Detroit, MI 48202, USA; Department of Medicine, University of California, San Francisco, San Francisco, CA 94143, USA; Center for Human Genomics and Personalized Medicine, Wake Forest School of Medicine, Winston-Salem, NC 27157, USA; Center for Research on Genomics & Global Health, National Institutes of Health, Bethesda, MD 20892, USA; Immunology Service, Universidade Federal da Bahia, Salvador, BA 401110170, Brazil; Department of Medicine, Northwestern University, Chicago, IL 60611, USA; Department of Medicine, University of Arizona College of Medicine, Tucson, AZ 85724, USA; Institute for Immunological Research, Universidad de Cartagena, Cartagena, 130000, Colombia; National Human Genome Center, Howard University College of Medicine, Washington, DC 20059, USA; Department of Microbiology, Howard University College of Medicine, Washington, DC 20059, USA; Caribbean Institute for Health Research, The University of the West Indies, Kingston, Jamaica; Departamento de Biorregulacao, Universidade Federal da Bahia, Brazil; Department of Medicine, Einstein Medical Center, Philadelphia, PA 19141, USA; Department of Neurology, University of California, San Francisco, San Francisco, CA 94158, USA; Institute for Human Genetics, University of California, San Francisco, San Francisco, CA 94143, USA; Department of Bioengineering and Therapeutic Sciences, University of California, San Francisco, San Francisco, CA 94143, USA; California Institute for Quantitative Biosciences, University of California, San Francisco, San Francisco, CA 94143, USA; Centro de Neumologia y Alergias, San Pedro Sula, 21102, Honduras; Facultad de Medicina, Universidad Catolica de Honduras, San Pedro Sula, 21102, Honduras; Department of Genetics and Genomics, Icahn School of Medicine at Mount Sinai, New York, NY 10029, USA; The Ann & Robert H. Lurie Children's Hospital of Chicago, Chicago, IL 60611, USA; Department of Pediatrics, Northwestern University, Chicago, IL 60611, USA; Department of Statistics, University of Chicago, Chicago, IL 60637, USA; Department of Medicine, University of Chicago, Chicago, IL 60637, USA; Institute for Genome Sciences, University of Maryland School of Medicine, Baltimore, MD 21201, USA; Program in Personalized and Genomic Medicine, University of Maryland School of Medicine, Baltimore, MD 21201, USA; Department of Medicine, University of Maryland School of Medicine, Baltimore, MD 21201, USA; Laboratório de Patologia Experimental, Centro de Pesquisas Gonçalo Moniz, Salvador, BA 4029-6710, Brazil; Department of Medicine and Center for Global Health, University of Chicago, Chicago, IL 60637, USA; Faculty of Medical Sciences, The University of the West Indies, Queen Elizabeth Hospital, Bridgetown, St. Michael, BB11000, Barbados; Department of Internal Medicine, Henry Ford Health System, Detroit, MI 48202, USA; Department of Physiology and Biophysics, University of Mississippi Medical Center, Jackson, MS 39216, USA; Department of Human Genetics, University of Chicago, Chicago, IL 60637, USA; Department of Epidemiology, Bloomberg School of Public Health, JHU, Baltimore, MD 21205, USA; Department of Biostatistics, Bloomberg School of Public Health, JHU, Baltimore, MD 21205, USA; Centre de Recherches Médicales de Lambaréné, 13901, Gabon; Institut für Tropenmedizin, Universität Tübingen, 72074, Germany; Department of Chemical Pathology, University of Ibadan, Ibadan, 900001, Nigeria; Department of Parasitology, Leiden University Medical Center, Netherlands; Faculty of Medicine, Universidad Nacional Autonoma de Honduras en el Valle de Sula, San Pedro Sula, 21102, Honduras; Department of Medicine, University of Mississippi Medical Center, Jackson, MS 39216, USA; Pulmonary and Critical Care Medicine, Morehouse School of Medicine, Atlanta, GA 30310, USA; Data and Statistical Sciences, AbbVie, North Chicago, IL 60064, USA; Illumina, Inc., San Diego, CA 92122, USA; Department of Medicine, Vanderbilt University, Nashville, TN 37232, USA; Department of Preventive Medicine, Northwestern University, Chicago, IL 60611, USA; CIBER de Enfermedades Respiratorias, Instituto de Salud Carlos III, Madrid, 28029, Spain; Departments of Biomedical Engineering and Biostatistics, Johns Hopkins University, Baltimore, MD 21205, USA; Department of Child Health, The University of the West Indies, Kingston, Jamaica; Biomedical Sciences Graduate Program, University of California, San Francisco, San Francisco, CA 94158, USA; Centro Medico de la Familia, San Pedro Sula, 21102, Honduras; Department of Pathology, Microbiology and Immunology, Vanderbilt University, Nashville, TN 37232, USA; Department of Parasitology, Leiden University Medical Center, Leiden, 2333, Netherlands

## Abstract

**BACKGROUND:** Asthma is a complex disease with striking disparities across racial and ethnic groups, which may be partly attributable to genetic factors. One of the main goals of the Consortium on Asthma among African-ancestry Populations in the Americas (CAAPA) is to discover genes conferring risk to asthma in populations of African descent.

**METHODS:** We performed a genome-wide meta-analysis of asthma across 11 CAAPA datasets (4,827 asthma cases and 5,397 controls), genotyped on the African Diaspora Power Chip (ADPC) and including existing GWAS array data. The genotype data were imputed up to a whole genome sequence reference panel from n=880 African ancestry individuals for a total of 61,904,576 SNPs. Statistical models appropriate to each study design were used to test for association, and results were combined using the weighted Z-score method. We also used admixture mapping as a complementary approach to identify loci involved in asthma pathogenesis in subjects of African ancestry.

**RESULTS:** SNPs rs787160 and rs17834780 on chromosome 2q22.3 were significantly associated with asthma (p=6.57 × 10^−9^ and 2.97 × 10^−8^, respectively). These SNPs lie in the intergenic region between the Rho GTPase Activating Protein 15 (*ARHGAP15*) and Glycosyltransferase Like Domain Containing 1 (*GTDC1*) genes. Four low frequency variants on chromosome 1q21.3, which may be involved in the “atopic march” and which are not polymorphic in Europeans, also showed evidence for association with asthma (1.18 ×10^−6^ ≤ p ≤ 3.06 ×10^−6^). SNP rs11264909 on chromosome 1q23.1, close to a region previously identified by the EVE asthma meta-analysis as having a putative African ancestry specific effect, only showed differences in counts in subjects homozygous for alleles of African ancestry. Admixture mapping also identified a significantly associated region on chromosome 6q23.2, which includes the Transcription Factor 21 (*TCF21*) gene, previously shown to be differentially expressed in bronchial tissues of asthmatics and non-asthmatics.

**CONCLUSIONS:** We have identified a number of novel asthma association signals warranting further investigation.

## BACKGROUND

Asthma is a complex disease where the interplay between genetic factors and environmental exposures controls susceptibility and disease progression. In the U.S., ethnic minorities are disproportionally affected by asthma. For example, African Americans and Puerto Ricans have higher asthma-related morbidity and mortality rates compared to European Americans [1-3]. In addition to environmental, cultural and socio-economic risk factors, genetic factors, possibly from a common background ancestry, likely underlie some of these disparities in the health burden of asthma in the U.S. Despite the relatively high burden of disease, representation of African ancestry populations in asthma genome-wide association studies (GWAS) to date has been limited. The largest asthma GWAS focused solely on African ancestry populations included only 908 asthma cases [4], and only 1,612 cases of African ancestry were represented in the largest asthma meta-analysis to date [5]. The Trans-National Asthma Genetic Consortium (TAGC) is currently conducting a meta-analysis of more than 100,000 subjects, of which only 2,149 are cases of African ancestry. The discovery of genetic risk factors for asthma in African ancestry populations has also been further hampered by legacy commercial genotyping arrays that do not provide adequate coverage of linkage disequilibrium (LD) patterns in African ancestry populations.

To address these research disparities and the paucity of information on African genetic diversity, we established the Consortium on Asthma among African-ancestry Populations in the Americas (CAAPA). We first performed whole genome sequencing (WGS) of samples collected from individuals (394 asthmatics, 424 non-asthmatics and 60 with unknown asthma status) who self-reported African ancestry from 19 North, Central and South American and Caribbean populations (446 individuals from 9 African American populations, 43 individuals from Central America, 121 individuals from 3 South American populations, and 197 individuals from 4 Caribbean populations), as well as individuals from continental West Africa (45 Yoruba-speaking individuals from Ibadan, Nigeria and 28 individuals from Gabon). These whole-genome sequences were made publicly available through dbGAP (accession code phs001123.v1.p1) and were incorporated into the reference panel on the Michigan imputation server (a free genotype imputation service, https://imputationserver.sph.umich.edu). The average sequencing depth of the CAAPA samples is ≈30x across the genome compared to the average depth of ≈7x in the 1000 Genomes Project (TGP) [6]. The CAAPA samples also represent 880 unique African genomes compared to 661 African TGP phase 3 genomes, thus making the CAAPA reference panel a valuable resource for imputing genetic variants of African origin.

Ongoing work in the CAAPA consortium has included coverage analysis of the novel variation identified in CAAPA sequencing. We found that only 69% of common SNP variants and 41% of low-frequency SNP variants identified by CAAPA can be tagged by traditional GWAS arrays at r^2^ >= .8.

In addition, lack of coverage of low frequency variants in GWAS arrays negatively impacts the imputation of low frequency variants. To address these issues, we used the CAAPA sequence data to develop the African Diaspora Power Chip (ADPC), a gene-centric SNP genotyping array designed to complement commercially available genome-wide chips, thereby improving tagging and coverage of African ancestry genetic variation [7]. The array included ~495,000 SNPs, with a minor allele frequency (MAF) that is enriched for low frequency variants (MAF between 0.01-0.05).

Using the ADPC, we genotyped CAAPA participants from nine sites (seven providing African American samples, one providing African Caribbean samples from Barbados and one providing samples from Puerto Rico), and then combined ADPC data with existing commercial genome-wide genotype data and imputed using the CAAPA reference panel. One additional African American dataset genotyped on the Affymetrix Axiom AFR [8] array was also imputed using the CAAPA reference panel.

We used these genotyped and imputed data to perform the largest GWAS of asthma in individuals of African ancestry to date (4,827 asthmatic cases and 5,397 controls).

## METHODS

### Participants and Disease Definitions

CAAPA investigators previously recruited African ancestry participants into asthma genetics studies across nine geographical sites (Table 1), most of which have been described elsewhere [9]. Participants in these studies were unrelated cases and controls, except for subjects from the Barbados Asthma Genetics Study (BAGS), where a family-based study design was used, and the Howard University Family Study (HUFS) where participants were a mixture of unrelated and related subjects. Definition of asthma was based on a doctor’s diagnosis and/or standardized questionnaires (see Supplementary Material for details); controls were defined as a reported negative history of asthma. The distributions of age, sex and age of asthma onset are summarized in Table S1.

**Table 1:**
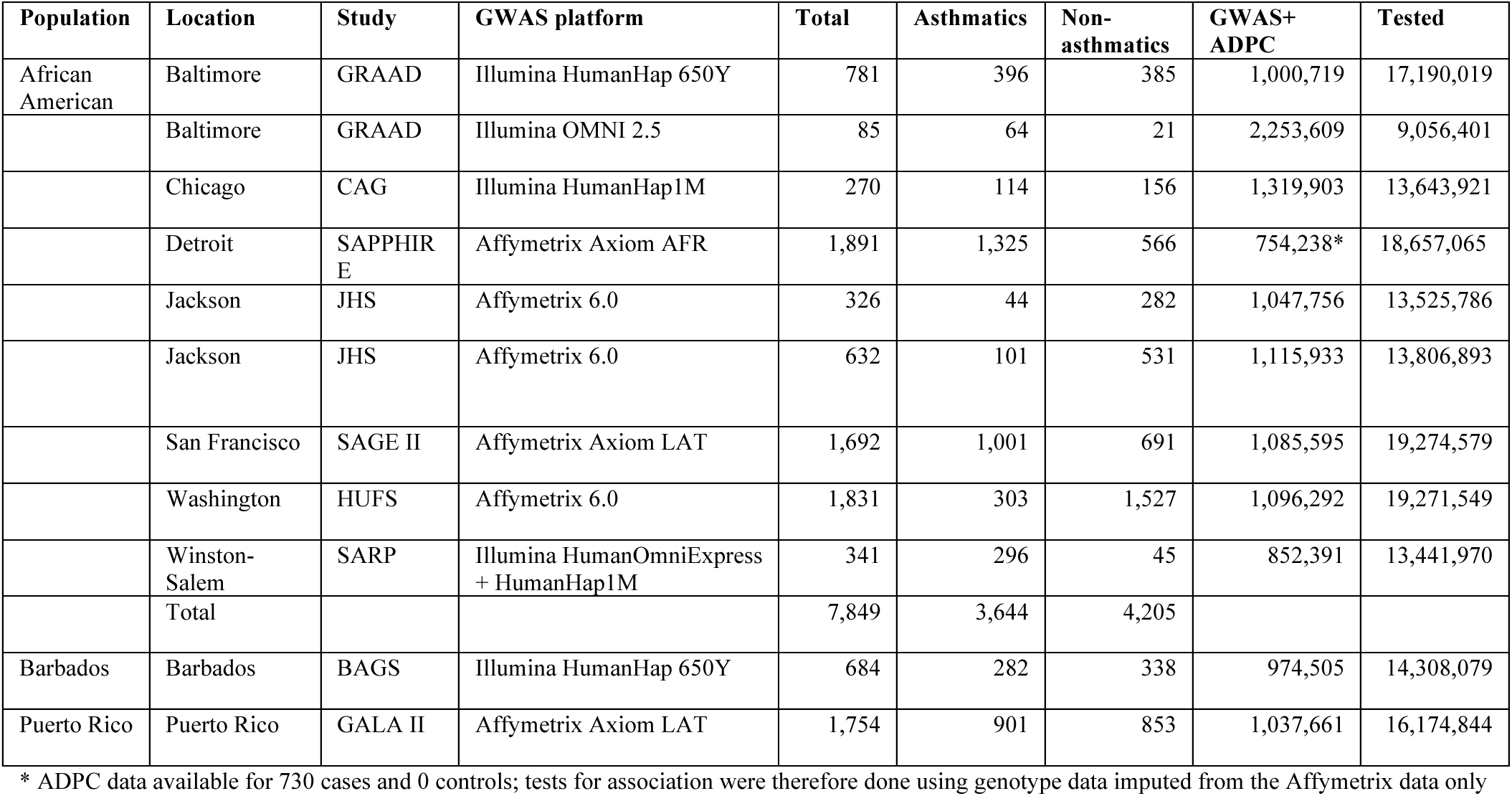
Studies included in the CAAPA meta-analysis. Studies were genotyped on a variety of GWAS backbones, and coverage of African genetic variation was augmented by additional genotyping on the African Diaspora Power Chip (ADPC). The table summarizes the final number of subjects included in the analysis per study, the number of genotyped SNPs in the combined GWAS backbone and ADPC array data sets, and the number of imputed SNPs that were used in the meta-analysis.

### Study Oversight

NIH guidelines for conducting human genetic research were followed. The Institutional Review Boards (IRB) of Johns Hopkins University (for GRAAD and BAGS), Howard University (for CRAD and HUFS), Wake Forest University (for SARP), the University of California, San Francisco (coordinating center for the SAGE II and GALA II studies), the Western Institutional Review Board for the recruitment in Puerto Rico (GALA II Puerto Ricans), Children's Hospital and Research Center Oakland and Kaiser Permanente-Vallejo Medical Center (for SAGE II), the University of Chicago (for CAG), University of the West Indies, Cave Hill Campus, Barbados (for BAGS), University of Mississippi Medical Center (for JHS), Henry Ford Health System (for SAPPHIRE), all reviewed and approved this study. All participants provided written informed consent.

### Genotyping

#### ADPC

Prior to genotyping on the ADPC, CAAPA samples were separately genotyped on a variety of GWAS platforms (Table 1). Each of the study sites was responsible for and performed their own genotype calling (see details in the Supplementary Material). To improve coverage of genetic variation specific to African ancestry populations, CAAPA subjects with existing GWAS data were genotyped on the ADPC at Illumina, Inc. FastTrack Services [7]. Genotypes from the ADPC were called following recommendations from Illumina Infinium Technical Notes implemented in Illumina’s GenomeStudio software (v2011.1) [10].

### Quality Control, Data Merging and Imputation

Initial sample and SNP quality filtering of the CAAPA GWAS data sets, including genomic coordinate transformation to hg19, was performed separately by each of the sites (see the Supplementary Material). A custom pipeline was then used to perform additional QC and to merge each of these GWAS data sets with corresponding ADPC genotype data (described in detail in the Supplementary Material). Finally, these combined data sets were used for imputation, a process where genotype posterior probabilities of untyped genetic markers are estimated by matching a set of phased typed markers to reference haplotype panels. SNP-based imputations were done on the Michigan imputation server (https://imputationserver.sph.umich.edu), separately for each dataset (see the Supplementary Material for more information). Imputing untyped markers in datasets initially typed on a variety of platforms allowed us to combine results for meta-analysis over all datasets. Imputed genetic data sets also provide a higher density of markers and a much larger number of low frequency and rare variants [11, 12].

### Genome-wide and local ancestry estimation

The populations included in the asthma association analyses are genetically admixed populations, deriving their origins from populations that were previously demographically isolated (African, European and Native American populations). To account for and leverage the population structure resulting from this admixture, we estimated both genome-wide and local ancestry for all CAAPA subjects.

To identify population outliers, a set of MAF and linkage-disequilibrium (LD) pruned SNPs genotyped across all CAAPA subjects was merged with an overlapping set of markers from unrelated European and African subjects from TGP phase 3 (84+84 CEU and YRI subjects) and unrelated Native American subjects (43 NAT) from Mao et al. [13]. This merged dataset was used to estimate principal components as well as proportions of African, European and Native American ancestry for each CAAPA subject. Principal component analysis was also performed separately by dataset to account for differences in population structure by case-control status. More detailed information can be found in the Supplementary Material (under the heading **Genome-wide ancestry analysis**).

When admixture occurs between different population groups, recombination events result in chromosomes that are a mosaic of blocks of ancestry derived from the different source populations (called local ancestry segments). Local ancestry inference is a statistical technique that use the probability of recombination events to determine the boundaries of each local ancestry segment, and differences in allele and haplotype frequencies between putative source populations to classify the source ancestry of all segments. The local ancestry of chromosomal segments from 7,416 African American and African Caribbean Barbados CAAPA subjects was inferred using the software program RFMix, with African and European TGP phase 3 subjects serving as reference panels [14]. We inferred 15,824 segments of local ancestry across the genomes of these 7,416 subjects. A more detailed description of this process can be found in the Supplementary Material (see **Local ancestry inference**).

### Functional annotation

Several steps were used to explore the potential functional relevance of associations observed: 1) The Genotype-Tissue Expression (GTEx) eQTL browser was used to determine if a SNP is an eQTL; 2) RegulomeDB was used to determine if a SNP may have a regulatory effect; and 3) ANNOVAR gene, filter and region based annotations were examined (Supplementary Table S2).

### Statistical Analysis

#### Association analysis

We first tested for association between imputed allele dosages and case-control status. The imputed allele dosage is defined as the expected number of copies of the variant allele each individual carries, and is a continuous number ranging between 0 and 2 (incorporating the uncertainty of the imputed genotypes). Due to different study objectives of participating studies, and differences in potentially relevant factors such as age of onset and diagnostic criteria (see the Supplementary Material), we performed tests for association separately for each dataset, and then combined the results in a meta-analysis, allowing for the assessment of association signals that may be particular to any given type of study design. Appropriate statistical tests were used for each type of dataset (see Supplementary Table S3 and the Supplementary Material; due to different effect size statistics, association test results were combined using the weighed Z-score method [15]). P-values < 5 × 10^−8^ were considered statistically significant.

#### Candidate gene analysis

In addition to the GWAS approach, we also explored associations with known asthma candidate genes in the meta-analysis results. Due to the relatively lower African contribution in the Puerto Rican (GALA II) study, this pipeline was applied twice: Once to the meta-analysis of all CAAPA datasets, and once to a meta-analysis excluding GALA II.

Asthma associations with p < 10^−5^ were selected from the GWAS catalog and Phenotype-Genotype Integrator (PheGenI) databases (http://www.genome.gov/gwastudies/ and http://www.ncbi.nlm.nih.gov/gap/phegenily, respectively). Gene regions were defined as 1) ±1MB from a gene for variants that fell within a single gene, 2) ±1MB from the variant and gene for variants that did not fall within the gene itself but for which the databases reported a gene, 3) ±1MB from a variant for variants with no reported gene, and 4) ±1MB from the first and last gene for SNPs falling within a region with many reported genes. For each identified gene region, imputed data from the SAGE II dataset (the largest African American dataset with unrelated subjects) was used to estimate the number of independent SNPs in each gene region (see the Supplementary Material for more information). The p-values of all SNPs within a candidate gene region were multiplied by the number of independent tests estimated for that region to give Bonferroni corrected p-values. Corrected p-values < 0.05 were considered statistically significant.

#### Admixture mapping

Admixture mapping identifies putative disease causing variants that occur more frequently on local ancestry segments inherited from a particular source population. We performed admixture mapping by using linear mixed models (implemented in the EMMAX software) to test for association between the number of copies of African ancestry at a local ancestry segment (0/1/2) and asthma case-control status. We fitted 15,824 models (one for each local ancestry segment) to the 6,686 African American and African Caribbean Barbados subjects for whom local ancestry was inferred and where both cases and controls were available. The models used a covariance matrix to account for relatedness between all individuals, as well as population structure, which accounts for global ancestry. The genotype dataset (e.g. GRAAD(1), GRAAD(2), CAG, etc.) was included as a fixed effect in these models. To correct for multiple testing, we estimated the number of effective tests using the method described by Gao et al. [16], and then applied a Bonferroni corrected significance threshold of 0.05/(262 effective tests)=1.9 × 10^−4^ See the Supplementary Material for a more detailed description of this analysis (heading **Admixture mapping**).

#### Stratifying association tests by local ancestry

A logistic regression interaction model was used to determine if an association differed by local ancestry (where case-control status ~ allelic_dosage + dataset + allelic_dosage^*^nr_copies_African_ancestry, with nr_copies_African_ancestry encoded as factor variable with values 0, 1 or 2, and the *glm()* function in the R package is used with first the binomial family link and then a logit link). SNPs yielding p-values <10^−5^ in either of the meta-analysis (i.e. including or excluding GALA II), or in the candidate gene analysis, were tested using this model. Kinship matrices were used to exclude closely related subjects from these association tests (i.e. those having kinship coefficient > 0.25 with one or more of the other subjects in the dataset). This analysis was restricted to only those datasets contributing to the meta-analysis (e.g. if a SNP was filtered out for a particular dataset due to low imputation quality, that dataset was excluded from the local ancestry interaction model). Models with interaction p-values < 0.05 (using the anova.glm() function in R, test=Chisq) were further examined by stratifying the dataset according to 0, 1 or 2 copies of African ancestry at that locus, and then using an additive allelic test for association within each stratum, testing for differences between case and control allele count.

## RESULTS

### Population Structure

Population structure was assessed by principal component analysis (PCA), as well as genome-wide estimates of proportions of ancestry deriving from specified reference populations (ADMIXTURE [17]). These results are summarized in Supplementary Figures S1 and S2. Three African American subjects were identified as PCA outliers, and were excluded from further association analysis. The median percentages of African ancestry in African American, Barbados and Puerto Rican subjects were 82.2, 90.9 and 18.14% respectively.

### Association Analysis

The individual dataset and meta-analysis QQ plots showed little evidence of systematic test statistic inflation (Supplementary Figure S3 and Figure 1). Table 2 summarizes association results from two meta-analysis: a meta-analysis of all the CAAPA GWAS datasets, and a meta-analysis excluding Puerto Rican (GALA II) subjects, due to their relatively higher European ancestry. SNPs are shown if they have p < 10^−6^ in either of the meta-analyses and/or were within an asthma candidate gene region and passed a multiple testing significance threshold for that gene region in either of the two meta-analyses. A detailed breakdown of the association results by dataset is shown in Supplementary Table S4, and the results of the candidate gene analysis are summarized in Supplementary Table S5. SNPs with MAF < 0.01 and with fewer than 8 datasets contributing to the meta-analysis are excluded from Table 2, but are included in Supplementary Table S4.

**Figure 1:**
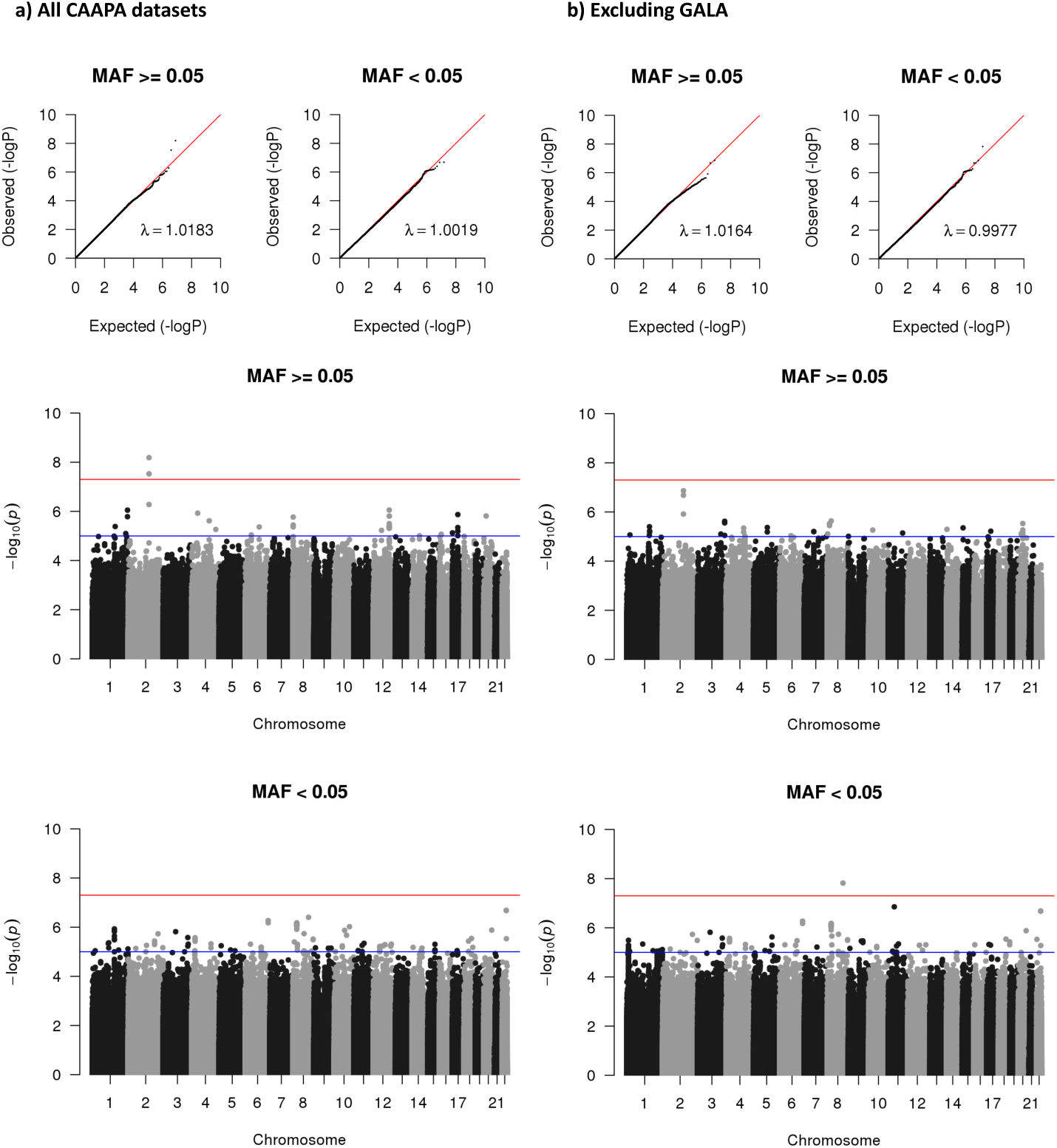
Summary of the distribution of meta-analysis p-values. QQ and Manhattan plots of the meta-analysis p-values, stratified by minor allele frequency (MAF) for low frequency and common SNPs. Inflation factors were calculated by transforming p-values to 1 degree of freedom (df) Chi-square statistics, and dividing the median of these statistics by the median of the theoretical Chi-square (1 df) distribution. The blue and red horizontal lines in the Manhattan plots represent the suggestive and significant p-value thresholds, respectively. Panel a) shows the QQ and Manhattan plots for the metaanalysis of all CAAPA datasets. Panel b) shows the QQ and Manhattan plots for the meta-analysis excluding GALA II.

**Table 2:**
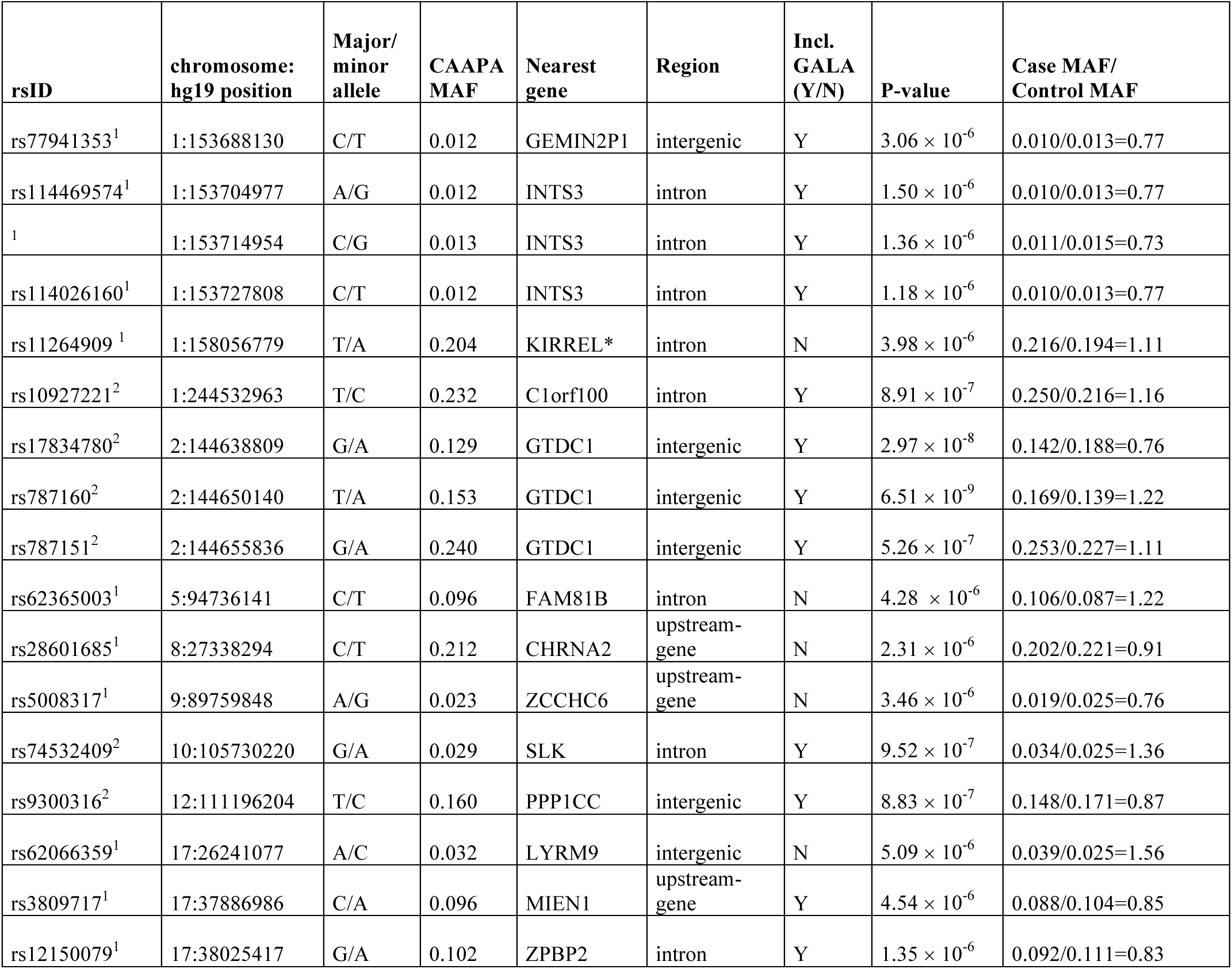
Summary of meta-analysis results. This table summarizes association results based on two meta-analysis: a meta-analysis of all the CAAPA GWAS datasets, and a meta-analysis that excludes GALA II, due to the relatively higher European ancestry within GALA II. The table summarizes association results that pass any of the following criteria: 1) SNPs with p-values < 10-6 in either of the meta-analyses that include or exclude GALA II, and 2) SNPs that fall within an asthma candidate gene region and that pass a multiple testing significance threshold for that gene region in either of the two meta-analyses. SNPs with MAF < 0.01 and with less than 8 studies contributing to the meta-analysis p-value are excluded from the table.

Two SNPs on chromosome 2q22.3, rs787160 and rs17834780, passed the genome-wide significance threshold of p < 5 × 10^−8^. These SNPs are intergenic and located close to the Glycosyltransferase Like Domain Containing 1 (*GTDC1*) and Rho GTPase Activating Protein 15 (*ARHGAP15*) genes (Supplementary Figure S4). *ARHGAP15* is expressed in lymph node cells, peripheral blood mononuclear cells, CD8 T-cells, and B-lymphocytes, while *GTDC1* is expressed in lung and peripheral blood leukocytes (genecards). Glycosyltransferase plays a role in the adhesion of environmental factors to epithelial cells, which could in turn play some role in risk to asthma [18]. SNP rs787160 falls in a region of weak transcription in fetal lung tissue (15 state model), and rs17834780 falls in a heterochromatin region in lung tissue (18 state model) in the Encyclopedia of DNA Elements (ENCODE) database.

Association between chromosome 17q21 and asthma are regarded as the most consistent finding in asthma genetics research to date [19]. Although SNPs in this region do not pass our genome-wide significant threshold in the meta-analysis results, two chromosome 17q21 SNPs, rs3809717 and rs12150079 (hg 19 base pair positions 37886986 and 38025417 respectively) were identified in the candidate gene analysis. SNP rs3809717 is upstream from the Migration and Invasion Enhancer 1 (*MIEN1*) gene and SNP rs12150079 is intronic to Zona Pellucida Binding Protein 2 (*ZPBP2*) gene. Interestingly, rs12150079 is annotated as a strong enhancer in the ENCODE B-lymphocyte and lymphoblastoid cell line (ENCODE 15 state model GM12878 cell line), and rs3809717 is annotated as either a strong or weak promoter in multiple ENCODE cell lines (ENCODE 15 state model, active promoter in the GM127878, HepG2, HSMM, HUVEC, NHEK and NHLF cell lines, weak promoter in the H1-hESC, HMEC and K562 cell lines).

A number of genes on chromosome 1q21.3 are putative asthma genes (*CRNN, LCE5A, CRCT1, LCE3E, IL6R, ADAR* and *KCNN3*). This region falls within the epidermal differentiation complex, which is also implicated in the development of atopic dermatitis [20]. Four SNPs falling within this region reached statistical significance in the candidate gene analysis, are in strong LD, and are within introns of the *INTS3* gene. These SNPs have a low MAF in the CAAPA datasets (MAF ~ 0.01, see Supplementary Table 4 for a more detailed breakdown), and are rare in European populations (MAF=0.001 in 1006 TGP phase 3 subjects). According to ENCODE, rs114469574 falls within enhancer regions in B-lymphocytes and lymphoblastoid cell lines, as well as the hepatocellular carcinoma cell line (15 state model, cell lines GM12787 and HepG2).

SNP rs11264909 on chromosome 1q23.1 reached significance in the candidate gene analysis that excluded GALA II. This SNP is located approximately 900 kb downstream from an association near the Pyrin and HIN Domain-Containing Protein 1 (*PYHIN1*) gene, which was reported as an African-ancestry specific association by the EVE asthma meta-analysis (rs1102000 at position 158,932,907 in the EVE meta-analysis, and rs1101999 at position 158,932,555 in the EVE replication; these 2 variants are in almost complete linkage disequilibrium in African Americans) [5]. SNP rs11264909 is intronic to the Kin of IRRE Like (Drosophila) (*KIRREL*) gene. Gene ontology (GO) annotations of *KIRREL* include myosin binding, which could affect bronchospasm [21]. ENCODE annotations for rs11264909 suggest this variant may affect transcription (the variant is annotated as having weak transcription in the H1-hESC cell line, transcription transition in HMEC and HSMM cell lines, insulator in the HUVEC cell line, and transcription elongation in NHEK and NHLF cell lines). The EVE SNPs rs1102000 and rs1101999 had p=2.58 × 10^−3^ and p=2.58 × 10^−3^ in the CAAPA meta-analysis when excluding GALA II, which is comparable to the replication p=2.80 × 10^−3^ reported by EVE for rs1101999 in African ancestry populations. The MAF of EVE SNPs rs1102000 and rs1101999 in the CAAPA datasets (excluding GALA II) ranged between 0.25 and 0.31, which is comparable to the MAF range of 0.26-0.29 reported by EVE. We note that the imputation quality metrics of the EVE SNPs rs1102000 and rs1101999 SNPs in the CAAPA datasets (Rsq ranging between 0.71-1.00, Supplementary Table 4) were on average slightly lower compared to the CAAPA SNP rs11264909 (Rsq ranging between 0.83-0.97, Supplementary Table 4). In addition, the majority of the CAAPA datasets did not have nominally significant p-values (i.e. < 0.05) in the individual datasets for these chromosome 1q23.1 SNPs (EVE SNPs rs1102000 and rs1101999 and CAAPA SNP rs11264909). For the CAAP SNP rs11264909, the direction of effect was consistent for 9 of the 10 datasets (with the smallest CAAPA dataset, GRAAD(2), having an opposite effect), contributing to an overall meta-analysis p=3.98 × 10^−6^. In contrast, for the EVE rs1102000 and rs1101999 SNPs, 3 datasets yielded conflicting effect directions (the JHS, GRAAD(2) and SARP datasets, see Supplementary Table S4).

### Admixture mapping

The distribution of p-values from the admixture mapping tests for association is summarized in Supplementary Figure S5. The QQ plot shows little evidence for systematic test statistic inflation. The deflated inflation factor of 0.90 appears to be largely driven by 0.1<p<0.5, whereas the number of p < 0.02 is greater than expected, suggesting that the association results are enriched with local ancestry segments showing differences in ancestry between cases and controls.

Only one segment of local ancestry, ranging from base pair positions 134,149,974-134,300,365 on chromosome 6, crossed the multiple testing threshold of p < 1.9 × 10^−4^ This segment falls within a local ancestry peak on chromosome 6q22.31-23.2, with increased African ancestry associated with increased risk of asthma (Supplementary Table S6). Genes falling in this segment include the Transcription Factor 21 (*TCF21*) and TATA-Box Binding Protein Like 1 (*TBPL1*) genes. The *TCF21* gene has been shown to be differentially expressed in bronchial biopsies of asthmatics compared to controls [22].

The second largest admixture-mapping peak occurred on chromosome 4q27-28.1. On average, asthma cases had higher African ancestry compared to controls in the local ancestry segments within this peak (Supplementary Table S6). The local ancestry segment with the smallest p-value ranged from base pair positions 124,387,497-124,567,967 and did not include any known genes. The closest gene to the segment is the Sprouty RTK Signaling Antagonist 1 (*SPRY1*) gene, which may negatively regulate respiratory organogenesis (genecards).

The admixture-mapping peak observed on chromosome 1q23.1 included both the *KIRREL* and *PYHIN1* genes (Supplementary Table S6), identified by the CAAPA and the EVE asthma meta-analysis, respectively. The effect of African ancestry is protective for all the local ancestry segments within this peak. Cases had on average 79.8% African ancestry and controls had on average 81.9% African ancestry in the segment with the smallest p-value.

### Stratifying association tests by local ancestry

SNPs with p < 10^−5^ in either of the meta-analyses (including vs. excluding GALA II), and SNPs associated with asthma in the candidate gene analysis, were tested for interaction with number of copies of African ancestry. Whereas admixture mapping identifies local ancestry segments that differ in ancestry between cases and controls, this approach stratifies the data by number of copies of African ancestry that an individual carries at each tested SNP. The interaction p-value indicates whether a difference in allele frequency between cases and controls is observed in one class of local ancestry background more than in another. Supplementary Table S7 summarizes these results; only two interactions reached significance, one specific to African ancestry, and one specific to European ancestry. The African specific interaction involved SNP rs11264909 on chromosome 1q23.1. A difference in allele frequency was observed only in those subjects carrying 2 copies of African ancestry in this region, with the minor A allele being observed more frequency in cases (0.12) compared to controls (0.10). For SNP rs10246878 on chromosome 7, allele frequency differences were only observed in carriers who had at least 1 copy of European ancestry.

## DISCUSSION

### Genome-wide association

Only two associations reached genome-wide significance in our study. The associations are found on chromosome 2q22.3, and to our knowledge, an association at this vicinity have only been reported once, in a GWAS of disocyanate asthma which included 74 cases and 824 controls of European ancestry [23]. Notably, the most significant SNP in our study, rs787160, is located one base pair from an insertion-deletion polymorphism. For this reason, the SNP was not included in the TPG phase 3 reference panel, which may explain why rs787160 has not been identified by any prior GWAS. The variant calling quality metrics of rs787160 in the CAAPA reference genomes did not indicate genotyping error (mean sequencing depth of 39.07 and mean genotyping quality of 89.54; see Supplementary Figure 6 for the distribution of these statistics). In addition, the imputation quality metrics in the CAAPA GWAS datasets were high (Rsq > 0.92 in all datasets, Supplementary Table S4). To obtain more evidence of the accuracy of this result, we genotyped the 3 most significant SNPs on chromosome 2q22.3 in HUFS samples using a TaqMan Real-Time PCR assay. In 1,792 subjects with at least one SNP to compare, the data was 98.76% concordant (Supplementary Table S8), which further substantiates that the SNPs were imputed with high accuracy. It is potentially interesting that SNP rs787160 was not identified by our stratified local ancestry models as having any African-specific effect and that the minor allele has a higher frequency in the GALA II dataset (i.e., especially among subjects having on average higher European ancestry, Supplementary Table S4). These observations suggest that the association is not specific to individuals with African ancestry. Although the associations with markers on chromosome 2q22.3 still need to be replicated in an independent sample set, our analysis underscores the importance of retaining SNPs adjacent to structural variants in reference panels, should they have variant call quality metrics of high quality.

### Candidate gene-analyses

The chromosome 17q21 association with asthma is evident in the meta-analysis including GALA II, but less evident when excluding GALA II (Supplementary Figure S4). This region has primarily been associated with childhood onset asthma [19, 24, 25]. In addition, we note that the two most significant associations with markers on chromosome 17q21 in the meta-analysis that included GALA II had a much higher MAF among Puerto Rican subjects (MAF 0.22 and 0.23 for rs3809717 and rs12150079, respectively) compared to subjects from the other CAAPA datasets (where MAF ranged between 0.05-0.08 for both rs3809717 and rs12150079, with the same minor alleles as in GALA II). Our models stratifying association tests by estimated local ancestry also failed to identify chromosome 17q21 SNPs as showing ancestry specific effects. The inclusion of adult onset asthmatics and reduced statistical power due to a lower MAF for SNPs in this region may explain the failure to achieve genome-wide significance for SNPs on chromosome 17q21 in our study.

Our candidate gene analyses identified a number of low frequency variants on chromosome 1q21.3 among African ancestry subjects that are virtually monomorphic among European ancestry populations. This region falls within the epidermal differentiation complex (EDC). The EDC region contains a very large and diverse family of genes associated with skin barrier dysfunction, and has been implicated in the development of atopic dermatitis (AD) [20]. Risk factors for AD are likely also involved in the pathogenesis of asthma, through the so-called “atopic march”, which suggests that allergic sensitization occurs first through a damaged skin barrier, and in turn affects the development of a person’s immune response and risk for asthma [26].

We identified a putative African-ancestry specific associated SNP, rs11264909, on chromosome 1q23.1. This variant is ~0.9MB downstream from 2 SNPs identified by the EVE asthma meta-analysis. Although the EVE SNPs were not polymorphic in Europeans, the A allele of the CAAPA rs11264909 SNP had a frequency of 0.20 in the CAAPA study populations (excluding GALA II) and a much higher frequency of 0.61 in Europeans (1006 TGP phase 3 subjects). The A allele was observed more frequently in CAAPA cases (frequency of 0.22, excluding GALA II) compared to CAAPA controls (frequency of 0.19, excluding GALA II), suggesting that cases on average have more European ancestry at this locus compared to controls. This was borne out by the admixture mapping results, which pointed to a protective effect of African ancestry in this region. These observations imply that the association observed with asthma is not “due to” African ancestry. However, when stratifying the association tests by the number of copies of African ancestry carried by each subject, only individuals homozygous for African ancestry at rs11264909 showed differences in allele frequency between cases and controls, with the A allele being observed more frequently among cases. This region may therefore indeed have an African ancestry specific effect, where the combination of African ancestry and the A allele at this locus increase the risk of asthma.

### Admixture mapping

The admixture mapping association analysis identified two regions, chromosomes 4q27-28.1 and 6q22.31-23.2, that were not immediately evident from the SNP association results. This corroborated the value of admixture mapping as a complementary approach to association mapping in admixed populations, which may uncover associations not detectable by SNP association tests alone [27]. The chromosome 6 local ancestry segment associated with asthma (after correcting for multiple tests) harbors the *TCF21* gene which is differentially expressed in the bronchial tissues of asthmatics and non-asthmatics [22]. Because no SAPPHIRE controls were typed on the ADPC, asthmatic cases from SAPPHIRE were not included in the admixture mapping tests for association. Consistent with the admixture mapping association results, the SAPPHIRE cases also have excess African ancestry in this local ancestry segment (>2 ± SD from the mean global African ancestry in SAPPHIRE cases), serving as independent verification that increased African ancestry may well increase risk for developing asthma. Furthermore, the SNP with the most significant association in this region, SNP rs111966851 (Supplementary Figure S4), has a MAF of 0.308 and 0.006 in Africans and Europeans respectively (based on 1322 African and 1006 European TGP phase 3 subjects), which also implies that African ancestry in this local ancestry segment increases risk for asthma.

### Functional relevance

We used a number of publicly available databases and resources, including GTEx [28], RegulomeDB [29] and ENCODE [30], to explore the potential functional relevance of the associations we identified in our study. We found no functional annotations with immediate or clear relevance to our peak SNPs. While these public resources have had tremendous value in extending our understanding of functional domains within the human genome, we acknowledge the following limitations that may have led to the lack of additional supporting findings for our peak SNPs: the lack of suitable target tissue of relevance in asthma (e.g. lack of peripheral blood mononuclear cells or nasal epithelium in public catalogs), the lack of the inclusion of diseased individuals in the public catalogs (i.e. none of the catalogs include asthmatics where the relevant regulatory signature may indeed be different) and perhaps most importantly the minimal representation of individuals with African ancestry in these public catalogs (i.e. all the added variation represented in our CAAPA genomes and imputed into the African ancestry case control subjects used in our approach is under-represented in the public catalogs and therefore potentially missed). Therefore, while our initial exploration leveraging these public catalogs yield no concrete evidence of functional relevance, we believe that the expansion of these catalogs over time will address some of our concerns.

## CONCLUSION

The prevalence, morbidity, and mortality related to asthma in ethnic groups with significant African ancestry, specifically Puerto Ricans and African Americans, is high compared to individuals of European descent and other Hispanic groups such as Mexican Americans. Thus, it is critical to understand the genetic architecture of asthma in these most afflicted subgroups of individuals with asthma. Because of recent admixture events over the past 500 years, these African descent populations have a complex genetic architecture. We accounted for this complex architecture through the development of our genotyping platforms, based on whole-genome sequencing of individuals of African descent with an overrepresentation of asthma. Our GWAS of >10,000 subjects of African ancestry represent among the largest studies to date, and demonstrates the complexity of asthma susceptibility genes in these recently admixed populations.

Our study has yielded SNPs and regions that may be specific to risk for asthma in individuals of African ancestry. In addition, our study provides evidence for a novel association on chromosome 2q22.3 that may be relevant in other ethnicities, which may have been undetected by previous asthma association studies due to poor representation of the associated variants in traditional imputation reference panels and genotyping arrays. The associations we report here warrant further investigation and replication. Further evaluation of the genes we identified in well-characterized asthma cohorts, already well represented within CAAPA, will provide additional insight into the role of these loci in determining asthma severity and progression. Another logical next step is the use of multi-omic approaches to generate hypotheses regarding their functional relevance, that is only limited now by the lack of multi-omic data sets that are representative of African ancestry.

## ACKNOWLEDGEMENTS

Goncalo Abecasis, Joshua Akey, Michael Bamshad, Jessica Chong, Aniket Desphande, Wenqing Fu, Rob Genuario, Wendy E. Grus, Lili Huang, Cindy Lawley, Devin P. Locke, Deborah Nickerson, Pat Oldewurtel, Romina Ortiz, Joseph Potee, Alexander Reiner.

## SUPPLEMENTARY MATERIAL

### CAAPA ADPC Participants and Disease Definitions

The primary selection criteria applied by each CAAPA study was (i) self-reported African ancestry; (ii) a physician's diagnosis of asthma or a negative history of asthma symptoms and asthma medication usage among controls; and (iii) the availability of DNA at a concentration of at least 10 ng/μl from a primary blood sample. 10,827 of the selected participants that had genotype data that passed all quality control criteria were used for the CAAPA association analysis (Table 1).

The **Genomic Research on Asthma in the African Diaspora (GRAAD) consortium** represent 8 studies of asthma in African American children and adults, and one additional study on healthy African Americans. GRAAD study subjects were recruited from the Baltimore-Washington DC metropolitan area, and all subjects used in this GWAS self-reported as African American. Case-control status was defined as previously described [1]. Briefly, standardized questionnaires administered by a clinical coordinator were used to determine whether a subject has a history of both self-reported and physician-diagnosed asthma (asthma case), or had no history of asthma (healthy control). Adult controls were favored to minimize the inclusion of subjects that may still develop asthma.

The **Chicago Asthma Genetics (CAG)** study comprises European American and African American families ascertained through affected sib pairs, case-parent trios (through affected offspring), adults and children with severe persistent asthma, and non-asthmatic control subjects (> 18 years) [2]. Asthma cases and families were recruited in the adult and/or pediatric asthma clinics at University of Chicago Hospital; controls were recruited from the medical center at large.

The **Study of Asthma Phenotypes and Pharmacogenomics Interactions by Race-ethnicity (SAPPHIRE)** is a longitudinal, population-based study aiming to identify the genetic predictors of asthma medication in a multiethnic patient [3]. Eligible patients received their care from a large healthcare system serving southeast Michigan and the greater Detroit metropolitan area. Patients with asthma were ≥ 12 years of age, had a documented physician diagnosis of asthma in the medical record, and had no prior diagnosis of congestive heart failure or chronic obstructive pulmonary disease [4, 5]. Control patients had the same enrollment criteria as case patients with the exception of having no prior diagnosis of asthma.

The **Jackson heart study (JHS)** was designed to study risk factors for of cardiovascular disease as well as other chronic disorders, such as asthma, in African American adults [6]. Detailed information on the study design and data collection has been published previously [7-9]. Participants were classified as asthmatic based on self-reported physician diagnosis of asthma or use of asthma medication [6].

The **Study of African Americans, Asthma, Genes & Environments (SAGE II)** is an ongoing population-based, case-control study recruiting African American participants from clinics in the San Francisco Bay Area [10]. Subjects were eligible if they were 8-21 years of age, self-identified all four grandparents as African Americans, and had <10 pack-years of smoking history. Asthma was defined by physician diagnosis and report of symptoms in the 2 years preceding enrollment. Controls had no reported history of asthma, eczema, hives, hay fever, allergic rhinitis, no reported use of medication for allergies and no reported symptoms of wheezing or shortness of breath during their lifetime

The **Howard University Family Study (HUFS)** is a population based family study of African Americans in the Washington metropolitan area. African American families, as well as unrelated individuals, were randomly ascertained to study the genetic and environmental basis of common complex diseases [11]. Asthma was defined based on self-reported physician diagnosis and medication history [12]. Controls were defined based on absence of physician-diagnosed asthma and no history of asthma medication.

The NHLBI-sponsored **Severe Asthma Research Program** (SARP1-2) is a multi-ethnic cohort of comprehensively characterized subjects with mild to severe asthma enriched for severe disease. SARP1-2 is an integrated study of the clinical and biological features of asthma with the goal of understanding the cellular and molecular mechanisms underlying asthma and its endotypes. Study participants that self-reported as African American were included in the CAAPA GWAS. A diagnosis of asthma in SARP was based on evidence of methacholine bronchial hyperresponsiveness or bronchodilator reversibility in combination with asthma symptoms [13].

Probands from the **Barbados Asthma Genetics Study (BAGS)** were recruited through referrals as described previously [1, 14, 15]. Nuclear and extended family members were also enrolled in the study. Asthma was defined based on a history of both self-reported and physician-diagnosed asthma, as well as a history of wheezing without an upper respiratory tract inflection [1]. Controls were selected based on no history of asthma.

The **Genes-environments & Admixture in Latino Americans (GALA II) is** an ongoing population-based, case-control study recruiting Latino children from five centers (Chicago, Illinois; Bronx, New York; Houston, Texas; San Francisco Bay Area, California; and Puerto Rico) [16]. Subjects were eligible if they were 8-21 years of age and all four grandparents self-identified as Latino. The definition of asthma cases and controls followed the same protocols as SAGE II. Only subjects recruited in Puerto Rico were included in the CAAPA GWAS.

### CAAPA WGS Reference Panel Participants and Disease Definitions

The whole genome sequence (WGS) reference panel that was used for genotype imputation is comprised of 880 unrelated subjects. Participant selection and disease definition is summarized in detail in Supplementary Appendix of Mathias et al. [17], but the note only describes the 642 subjects in the data freeze used for that study’s analysis. The final data freeze of 880 subjects used for genotype imputation in our association study is summarized in Supplementary Table S9. Samples collected from JHS, Gabon and Palenque were not available in the previous data freeze (642 subjects), but are available in the final data freeze (880 subjects). The JHS is described above (under **CAAPA ADPC Participants and Disease Definitions**). Asthma was not defined for the subjects from Gabon and Palenque.

### GWAS backbone genotyping, SNP calling and quality control

CAAPA studies were genotyped on a variety of GWAS backbones (Table 1). Each of the study sites was responsible for and performed their own genotype calling and genomic coordinate transformation where applicable, and for most datasets, this has been described previously (GRAAD(1)+BAGS: [1], CAG: [18], SAPPHIRE: [3], JHS: [19], SAGE II: [20], HUFS: [21], SARP [], GALA II [16]). Exceptions are described below.

#### GRAAD(2)

Samples from GRAAD participants that were recruited after the initial GWAS study[1] were genotyped on the Illumina Omni2.5 BeadChip at the Lowe Family Genomics core, Johns Hopkins. Genotypes were called following recommendations from Illumina Infinium Technical Notes implemented in Illumina’s GenomeStudio software (v2011.1).

SNPs that were out of HWE (p-value < 10^−6^) and SNPs with a missing call rate > 0.1 were dropped from the dataset. Samples with a missing call rate > 0.01 and samples with gender discrepancies (using F-statistics estimated by PLINK, samples with F(inbreeding males)<0.2 or F(inbreeding females)>0.5 defined as miss-classified) were dropped from the dataset. PLINK was used to estimate IBD between all pairs of subjects, using a set of LD-pruned SNPs. Duplicates (PLINK PI_HAT > 0.9, remove both individuals) and related individuals (PLINK PI_HAT > 0.35, remove at least one individual) were then dropped from the dataset.

#### JHS

After receiving the JHS datasets from the study coordinators, genomic coordinates were translated using CrossMap (http://crossmap.sourceforge.net) and the hg18tohg19 chain file published with this software. Genotyping on the Affymetrix 6.0 array was performed in two batches, and these two genotype data batches were therefore imputed and analyzed separately.

#### SARP

Genotypes were called using the GenomeStudio clustering algorithm. DNA samples with call rates <0.90 were excluded. SNPs with call frequencies <0.90 were excluded and clustering was repeated followed by exclusion of samples with a call rate <0.98. After updating SNP statistics, a second round of SNP exclusion was performed for cluster separation scores <0.20 followed by SNPs with score <0.30 and a GenTrain score <0.75. After clustering and genotype calling, QC was performed using PLINK. SNPs with a MAF >0.01 or with HWE p<0.001 in unrelated subjects were removed. Sex was determined for all subjects using chromosome X data and subjects with a mismatch with self-reported sex data were excluded. IBD tests were also performed to exclude identical or related samples.

### ADPC genotyping, SNP calling and quality control

10,411 CAAPA samples, 346 HapMap controls and 217 internal samples were genotyped at Illumina on the ADPC. 306 samples failed QC at Illumina. For the remaining 9,888 CAAPA samples, sample quality control was performed in the following order:

1. Remove IBD duplicates (n=193; performed on 113,324 LD-pruned SNPs filtered for SNPs that have Hardy-Weinberg equilibrium (HWE) p-values > 1e-6 and MAF > 0.05; samples with PLINK PI_HAT < 0.9 were classified as IBD duplicates)
2. Remove samples with a high proportion of missing genotypes (n=0, more than 1% missing genotypes)
3. Remove samples with gender discrepancies (n=28, using F-statistics estimated by PLINK, samples with F(inbreeding males)<0.2 or F(inbreeding females)>0.5 were removed)
4. Remove samples with excess heterozygosity (n=2, using F-statistics estimated by PLINK, samples with F(inbreeding)>0.25).

An additional 364 samples without existing GWAS data, and 71 samples from two studies too small for accurate imputation (the imputation server requires at least 50 samples) were also removed, leaving 9230 samples.

Preliminary SNP QC was performed using GenomeStudio. 130,637 SNPs that failed one or more of the following criteria were removed, as recommended by Illumina:

1. Call frequency < 0.97
2. Replicate errors > 2
3. Parent-parent-child (PPC) errors > 2
4. Cluster separation <= 0.3
5. Mean normalized intensity of first homozygote cluster (AA_R_Mean) <= 0.2 and/or mean normalized intensity of heterozygote cluster (AB_R_Mean) <= 0.2 and/or mean normalized intensity of second homozygote cluster (BB_R_Mean) <=0.2
6. Mean of normalized theta value for first homozygote cluster (AA_T_Mean) > 0.3
7. Standard deviation of normalized theta value for first homozygote cluster (AA_D_Mean) > 0.06
8. Mean of normalized theta value for second homozygote cluster (BB_T_Mean) < 0.7
9. Standard deviation of normalized theta value for second homozygote cluster (BB_D_Mean) > 0.06.

### Merging and imputation of GWAS and ADPC datasets

After receiving GWAS SNP array data from the various study centers, a pipeline of custom scripts was used to apply consistent quality control (QC) metrics to the GWAS datasets and to merge autosomal content from the GWAS and ADPC datasets. The pipeline was applied separately to each of the GWAS datasets listed in Table 1, and each of the datasets were uploaded separately to the imputation server hosted by the University of Michigan (https://imputationserver.sph.umich.edu; for the imputation configuration, the CAAPA - African American reference panel was used, the configured population was AA (CAAPA), and ShapeIT was used for phasing).

The steps applied in the pipeline were as follows:

1. Non-autosomal SNPs and SNPs with duplicate positions were removed from the GWAS dataset. Samples that were not typed successfully on the ADPC array were also removed. SNPs that failed a test for Hardy-Weinberg equilibrium (HWE; p-value < 0.0001), monomorphic SNPs and SNPs that had more than 5% missing genotypes were then removed from the GWAS dataset.
2. Non-autosomal SNPs and four SNPs with duplicate positions were removed from the ADPC dataset. SNPs that failed a test for Hardy-Weinberg equilibrium (p-value < 0.0001), monomorphic SNPs and SNPs that had more than 5% missing genotypes were removed.
3. AT/CG SNPs were handled as follows, in both the GWAS and ADPC datasets: SNPs not in the reference panel were deleted; SNPs with MAF > 0.4 in either the dataset or reference panel were deleted; SNPs that differ in MAF more than 1% compared to the reference panel were deleted; for the remaining SNPs, if the minor allele was different from the reference panel minor allele, the strand of the SNP was flipped. These thresholds are stringent and a large number of SNPs were deleted (roughly 30 000 SNPs on the ADPC); however this strategy was preferred as incorrect AT/CG SNPs could potentially have a severe impact on imputation quality.
4. Next, the GWAS and ADPC datasets were uploaded to the imputation server for pre-imputation QC (Quality Control Only option). Based on the QC report downloaded from the server, SNPs typed on alternate strands compared to the reference panel were identified, and the alleles of these SNPs were flipped.
5. SNPs with common positions in the GWAS and ADPC datasets were extracted in order to identify discordant samples, and to verify the genotype concordance between the datasets. The proportion of alleles shared identical by-descent (IBD) between all pairs of samples were estimated using PLINK (PI_HAT output column), and GWAS and ADPC samples with identical sample identifiers but sharing less than 0.9 alleles IBD were removed from the GWAS and ADPC datasets (21 samples). The concordance between the remaining GWAS and ADPC samples was then checked (concordance was > 99.7% in all datasets).
6. Next, genotypes for corresponding GWAS and ADPC samples were merged into a single file. Discordant SNPs were handled as follows (same SNP in the ADPC and GWAS datasets, for the same sample, with different genotype calls): if a SNP was discordant in >= 5% of samples, the SNP was deleted from the dataset, else the SNP value was set to missing for the sample(s) it was discordant for.
7. Merged genotype files were uploaded to the imputation server. Imputed datasets and reports were downloaded after the job runs completed.
8. Based on the findings of a study that quantified imputation accuracy in African Americans, post-imputation SNP filtering was done based on minor allele frequency (MAF) and the per-SNP estimation of the squared correlation between imputed allele dosages and true unknown genotypes (Rsq) [22]. SNPs with MAF ≤ 0.005 were excluded if Rsq ≤ 0.5 and SNPs with MAF > 0.005 were excluded if Rsq ≤ 0.3.

The above process is depicted in Supplementary Figure S7 for the HUFS dataset. The pipeline automatically generated flow diagrams and the diagrams were manually checked for each dataset to verify the correctness of each step in the process. Results for the data merging steps and results scraped from reports downloaded from the imputation server are summarized in Supplementary Table S10.

### Imputation pipeline

#### Creating the reference panel

1. **Subject removal:** Identity-by-descent (IBD) tests were performed on 917 CAAPA samples to test for unexpected relatedness. IBD tests were generated from the multi-sample VCF by KNOME, Inc. Subjects were removed in order to ensure that no duplicate, parent-offspring or sibling relationships are present in the data.
2. **Variant filtering:** Variants with genotyping quality scores (GQ) less than 20, depth (DP) less than 7, and in regions of segmental duplication were removed. Only bi-allelic variants with call rates of at least 95% were retained.
3. **Phasing:** Phasing was performed using ShapelT (v2.r790) using 0.5 Mb windows.

As part of collaboration with the University of Michigan, this phased reference panel was uploaded to the Michigan imputation server.

#### Imputation

Full details of the pipeline is documented at

https://imputationserver.sph.umich.edu/index.html#!pages/pipeline. Briefly, the steps in the pipeline is as follows:

1. **Create chunks** of size 20 Mb.
2. **Do quality checks on chunks.** For chunks, the amount of valid variants, the amount of variants found in the reference panel, and the sample call rate are checked. For variants, valid alleles (A/C/G/T), SNP call rate, allele frequency differences between study and reference panel, and allele switches and strand flips are checked.
3. **Phase genotypes.** 3 phasing algorithms are available: ShapeIt, HapiUR and Eagle. For this study, we used ShapeIt to phase genotypes.
4. **Impute genotypes per chunk** using Minimac3 and a window size of 500 Kb.
5. **Merge chunks into a single file, per chromosome.**

### Quantifying imputation accuracy

Post-imputation SNP filtering was done based on MAF and the per-SNP Rsq [11]. On average, 5.5% SNPs with MAF > 0.005 were dropped from each study data set. Excluding the GRAAD(2) dataset, 72.6% of SNPs with MAF ≤ 0.005 were deleted (Supplementary Table S11). In contrast, 96% of GRAAD(2) SNPs with MAF ≤ 0.005 were dropped, most likely due to the small size of this sample (88 individuals), which may have affected the accuracy of the pre-phasing step performed during imputation. (Note: a MAF of 0.005 translates to the occurrence of a rare allele in 1 out of 100 individuals; imputation of allelic dosages rather than integer allele counts means it is still possible to observe a rare allele in only 88 individuals). A summary of the median Rsq values by chromosome and dataset is shown in Supplementary Figure S8. Datasets with the largest density of SNPs performed best overall (i.e. the GRAAD(2) and CAG datasets had the highest Rsq values, despite their small sample sizes). Similar to the results of Liu et al., imputation quality was lowest for chromosome 19, which may be due to the high gene density of this chromosome [11].

To further gauge imputation accuracy, 15% of the overlapping SNPs genotyped in the GRAAD(2), HUFS and GALA II datasets on commercial arrays (i.e. not on the ADPC) were selected at random and omitted from the combined datasets (10,751 SNPs in total that are also present in the CAAPA reference panel), and the imputations were run again. In terms of sample size, the first two datasets are the smallest and largest African American sample sizes respectively, and the GRAAD(2) dataset had the highest density of genotyped SNPs. GALA II participants are from Puerto Rico, and this dataset was examined in detail due to differences in the population structure of Puerto Rico compared to the other studies (Supplementary Figures S1 and S2). Imputation error can be quantified by comparing typed SNPs and most probable imputed genotype at these masked SNPs. This comparison was done per SNP, yielding an accuracy proportion per SNP (number of imputed genotypes that matched the typed genotypes exactly, divided by the number of total genotypes that were compared). The median and interquartile range accuracies (i.e. the proportion of matching imputed and typed genotypes for a particular SNP) for GRAAD(2), GALA II and HUFS were 0.9886 [0.9545 – 1.0000], 0.9618 [0.9236 – 0.9829] and 0.9610 [0.9194 – 0.9838], respectively, indicating high imputation accuracy.

### Genome-wide ancestry analysis

Genome-wide ancestry of subjects was assessed in order to identify population outliers to exclude from association analysis, as well as gauging the degree of population structure in the various studies (differences in ancestry between cases and controls).

#### Methods

First, SNPs with MAF < 1% were removed, followed by a linkage-disequilibrium (LD) pruning step. These steps were applied separately for each of the three ethnicities represented by the CAAPA studies (African American, African Caribbean = Barbados and Puerto Rican), and the datasets were then restricted to pruned SNPs common to all (122,206 SNPs). The algorithm implemented in the PLINK software package was used for the LD pruning step; SNPs within a window were recursively removed using a variance inflation factor threshold (window size 50 SNPs, step size 5 SNPs, variance inflation factor threshold 2). Note that for the SAPPHIRE dataset, the combined genome-wide ancestry analysis described here was restricted to the 730 asthmatic cases for which ADPC data were available.

Unrelated phase 3 thousand genomes project (TGP) subjects of European ancestry (CEU, n=84) and African ancestry (YRI, n=84), as well as unrelated Native American (NAT) subjects from Mao et al. [18] (n=43), were used as reference panels for the genome-wide ancestry analysis. Using the combined dataset of SNPs common to the reference panels plus the pruned CAAPA SNPs (16,410 SNPs), principal components were formed from the genotypes of the CAAPA and reference subjects by the R Bioconductor package. This software package calculates principal components by partitioning subjects into related and unrelated subsets (where the partitioning is based on a kinship matrix, see **Genetic covariance** structure). PC1 distinguished African ancestry from European and Native American ancestry and PC2 distinguished Native American ancestry from European and African ancestry. Based on this analysis, three outlier individuals that clustered close to the TGP CEU (European) subjects were identified, and these subjects were removed from the datasets prior to association testing.

The combined dataset (16,410 SNPs) was also used to estimate the proportion of genome-wide ancestry deriving from the three source populations represented by the reference panels, for each CAAPA subject. This was done using the software program ADMIXTURE. Because the BAGS and HUFS studies include families, and ADMIXTURE assumes that subjects are unrelated, only the founders in these studies (n=226 and n=997 respectively) were included for the ADMIXTURE analysis. The results are summarized in Supplementary Figure S2 and Supplementary Table S12.

Finally, the GENESIS PCA was repeated separately for each of the CAAPA datasets using the CAAPA pruned SNPs (122,206 SNPs), in order to test for differences in ancestry between cases and controls within each dataset. Tests for association between the top 10 principal components and case-control status were done separately for each principal component using logistic regression (*glm()* function in the R *base* package, binomial family with a logit link). For the JHS and GALA II study participants, association between ADMIXTURE estimates of genome-wide ancestry and case-control status was also tested using logistic regression (*glm()* function in the R *base* package, binomial family with a logit link), separately for each study and ancestry.

#### Results

The first principal component explained most of the variance in the genetic data (Supplementary Figure S9) in all of the datasets, which was similar to previous results in a GWAS of asthma in African ancestry populations [1]. Furthermore, differences in genome-wide ancestry between cases and controls in majority African ancestry populations were minimal. These differences were only evident on the first principal component in 2 of the 9 African American datasets, and were not detectable in Barbados. Only CAG, SAPPHIRE and HUFS had lower order principal components associated with asthma status (Supplementary Table S12). Differences in genome-wide ancestry were however observed between African American cases and controls from JHS (as evident on principal component 1, see Supplementary Table S12), and the Puerto Rican cases and controls from GALA II (as evident on principal components 1-3, see Supplementary Table S12). For these 3 datasets, cases had on average slightly, yet statistically significant, higher proportions of African ancestry compared to controls (Supplementary Figure S10).

### Association analysis

Linear mixed effects models were used for association testing of the CAG, GRAAD, GALA II and SAGE datasets (as implemented in the EMMAX software package). This model uses a covariance matrix to account for relatedness between all individuals, as well as the observed population structure. This linear model has been shown to yield a good approximation for significance testing in case-control GWAS, when the number of cases and controls in a dataset is balanced [23]. However, the approximation breaks down for an unbalanced number of cases and controls, which was the case for some of the CAAPA datasets. For this reason, logistic regression was used to test for association in the JHS(1), JHS(2), GRAAD(2), SAPPHIRE and SARP datasets (as implemented in the PLINK software package), after ensuring the coefficient of relatedness between each pair of subjects was below 0.25. The first principal component, as well as any of the top 10 principal components associated with asthma status (p < 0.05, Supplementary S12), were included as covariates in these logistic regression models. Unrelated and family-based designs were used to recruit the HUFS subjects, and although this dataset has an unbalanced number of cases and controls, due to the high degree of relatedness between family members, fixed effects only regression models are not suited for association analysis in this dataset. Instead, logistic (and not linear) mixed effects models were used to test for association (as implemented in the GENESIS R Bioconductor package). This test uses a kinship matrix to encapsulate relatedness and excludes other sources of variance such as population structure, and therefore principal components were also included as fixed effect covariates, using the same inclusion strategy described above for the logistic regression models implemented in PLINK. The BAGS cohort is also comprised of families ascertained through asthmatic cases, and includes 64 subjects with unknown phenotypes. To maximize statistical power, a quasilikelihood score test was used to analyze BAGS families (as implemented in the MQLS software package) [24], using a specified disease prevalence of 0.2 [25]. This test allows both linkage and association to contribute to the test statistic, and also leverages genetic information from subjects with unknown phenotypes.

Additional filtering of the association results was required prior to performing the meta-analysis. A more stringent Rsq filter was applied to the BAGS data to remove spurious p-values (Supplementary Figure S11). In addition, based on recommendations from the GIANT consortium [26], SNPs with a minor allele count (MAC) ≤ 6 were removed, which diminished test statistic inflation initially observed in some of these CAAPA datasets (Supplementary Figure S12). After performing these steps, the individual dataset and meta-analysis QQ plots showed little evidence of test statistic inflation (Supplementary Figure S3 and Figure 1).

### Candidate gene analysis

Asthma associations with p < 10^−5^ were selected from the GWAS catalog and Phenotype-Genotype Integrator (PheGenI) databases (http://www.genome.gov/gwastudies/ and http://www.ncbi.nlm.nih.gov/gap/phegenily, respectively). Gene regions were defined as 1) ±1MB from a gene for variants that fell within a single gene, 2) ±1MB from the variant and gene for variants reported for a gene but did not fall within the gene itself, 3) ±1MB from a variant for variants with no reported gene, and 4) ±1MB from the first and last gene for SNPs falling within a multi-genic region. Gene boundaries were based on USCS coordinates, and for those genes not in USCS, based on ENSEMBL BIOMART coordinates, and for those genes not in USCS or ENSEMBL BIOMART, based on the NCBI gene database coordinates.

Imputed data from the SAGE II dataset (the largest African American dataset with unrelated subjects) was used to estimate the number of independent SNPs in each identified candidate gene region. This was done using PLINK’s sliding window LD-prune algorithm (window size of 50 SNPs, step size of 5 SNPs, and a variance inflation factor threshold of 5, corresponding to a multiple correlation coefficient of ≤ 0.8 between SNPs). The number of SNPs selected by the PLINK algorithm for a region was used as a proxy for the number of independent tests in that region. The p-values of all SNPs within a candidate gene region were multiplied by the number of independent tests estimated for that region to give Bonferroni corrected p-values. Corrected p-values < 0.05 were considered statistically significant.

### Genetic covariance structure

Kinship matrices were estimated employing the same set of 122,206 CAAPA pruned SNPs used for genome-wide ancestry analysis.

Balding-Nichols kinship matrices were generated using the EMMAX software package and were included in the linear mixed effect models used to test for association between SNPs and asthma (CAG, GRAAD, BAGS, GALA II and SAGE; EMMAX software), as well as for the admixture mapping association tests. The matrices were used to account for relatedness between individuals as well as population structure (differences in ancestry between cases and controls).

Kinship matrices were also estimated using the KING-robust algorithm implemented in the KING software package [27], and was specified in the GENESIS logistic mixed effects models used for association tests of HUFS. The matrices are robust to the presence of population structure, and the KING kinship matrices were therefore also estimated and used to identify pairs of related subjects to exclude from logistic regression association tests. For each pair of subjects that shared a kinship coefficient > 0.25 (KING Kinship column × 2), the subject with the least represented phenotype were deleted (controls for GRAAD(2), SAPPHIRE, JHS(1), JHS(2), cases for SARP); if both subjects had the same phenotype the sample with the highest missing genotype rate in the ADPC dataset were deleted. 2 controls and 1 case, 1 control, 15 controls, and 6 cases were deleted from the GRAAD(2), JHS(1), JHS(2) and SARP datasets respectively.

### Local ancestry inference

First, GWAS array and ADPC genotyped autosomal SNPs that overlapped in 7,416 African American and Barbados subjects were merged in order to perform haplotype phasing using the ShapeIT software package (361,411 SNPs). After performing phasing, the 7,416 genomes were merged with African and European TGP phase 3 genomes (99 CEU and 108 YRI subjects, files downloaded from

https://ftp.1000genomes.ebi.ac.uk/vol1/ftp/release/20130502/to_data/raw/tgp_release_20130502). The genetic map required as input for local ancestry inference was downloaded from https://github.com/joepickrell/1000-genomes-genetic-maps/tree/master/interpolated_from_hapmap. Custom scripts were used to create input files for local ancestry inference according to the RFMix input file specification. The final phased input files comprised 273,738 markers.

RFMix inferred 15,824 local ancestry segments across the CAAPA genomes. The mean proportion of African ancestry inferred for each of these segments is summarized graphically in Supplementary Figure S13. Deviations between mean local African ancestry and mean genome-wide African ancestry are normally distributed, as expected (Supplementary Figure S14). Estimates of genome-wide ancestry from ADMIXTURE and RFMix were also highly concordant (Supplementary Figure S15), with a mean estimate difference of only 0.0009 (less than 0.1%).

### Admixture mapping

Custom scripts were used to convert the RFMix local ancestry calls to an EMMAX dosage TPED file, where the encoded “dosage” is defined as having 0, 1 or 2 copies of African ancestry at a particular local ancestry segment. Tests for association between the local ancestry dosage values and asthma case-control status were then done using the linear mixed effect models implemented in the EMMAX software package. Phenotypes for subjects from SAPPHIRE, which included only asthmatics and no controls on the ADPC, as well as the 3 population outliers identified by the genome-wide ancestry analysis, were set to missing. The models included dataset as a fixed effect covariate, and a Balding-Nichols kinship matrix as random effect.

The method described by Gao et al. was used to estimate the number of effective tests that should be used in a Bonferroni correction for multiple testing [28]. Briefly, a local ancestry correlation matrix, and corresponding eigenvalues, was calculated using local ancestry dosage values (0/1/2 copies of African ancestry) per chromosome. The number of effective tests for a chromosome is then set to the number of eigenvectors that explain 99.5% of the variance in the local ancestry data (for n local ancestry segments, find the largest k such that ∑^*k*^_*i*=1_ *eigenvalues* ∑^*n*^_*i*=1_ *eigenvalues* ≤ 0.995). The total number of effective tests is then set to the sum of the number of effective tests calculated for each of the 22 autosomes. Using this method, a Bonferroni corrected p-value threshold of 0.05/(262 total number of effective tests)= 1.9 × 10^−4^ was used to claim statistical significance.

### TaqMan genotyping and imputation data concordance

Genotyping was performed on the ABI 7900HT Sequence Detection System on 3 single nucleotide polymorphisms (SNPs). Using samples from HUFS, the top three SNP associations on chromosome 2q22.3 were genotyped in order to check for concordance with imputed genotypes (rs17834780, rs787151 and rs787151).

#### SNP Genotyping

1. Genomic DNA was extracted from peripheral blood samples.
2. Genotyping was performed using the TaqMan™ probe based, 5' nuclease assay with minor groove binder (MGB) chemistry (ABI, Foster City,CA). Genotyping was performed at a total volume of 5 μL per well, in 384- well MicroAmp Optical plates, using real-time detection followed by an endpoint read (Allelic discrimination) using the ABI 7900 HT Sequence Detection System software SDS2.4 (Applied Biosystems, Foster City, CA, USA).
3. Fluorescence data files from each plate were analyzed on scatter plots in the allelic discrimination window by automated allele calling software SDS 2.4 (ABI, Foster City, CA) and visually inspected.
4. Quality control: 5 – 10% of the randomly selected samples were repeated as routine quality control procedure.

#### Genotype concordance

TaqMan genotype call quality was assessed by means of manual inspection of the allelic discrimination plots, checking the proportion of successful genotype calls, and tests for HWE. The imputed genotype calls for corresponding samples were then converted to “hard” genotype calls, for those genotypes with imputation probabilities of at least 80%. The percentage of matching TaqMan and imputed genotype calls was then calculated. These results are summarized in Supplementary Table S8.

### SUPPLEMENTAL FIGURES

**Supplementary Figure S1:**
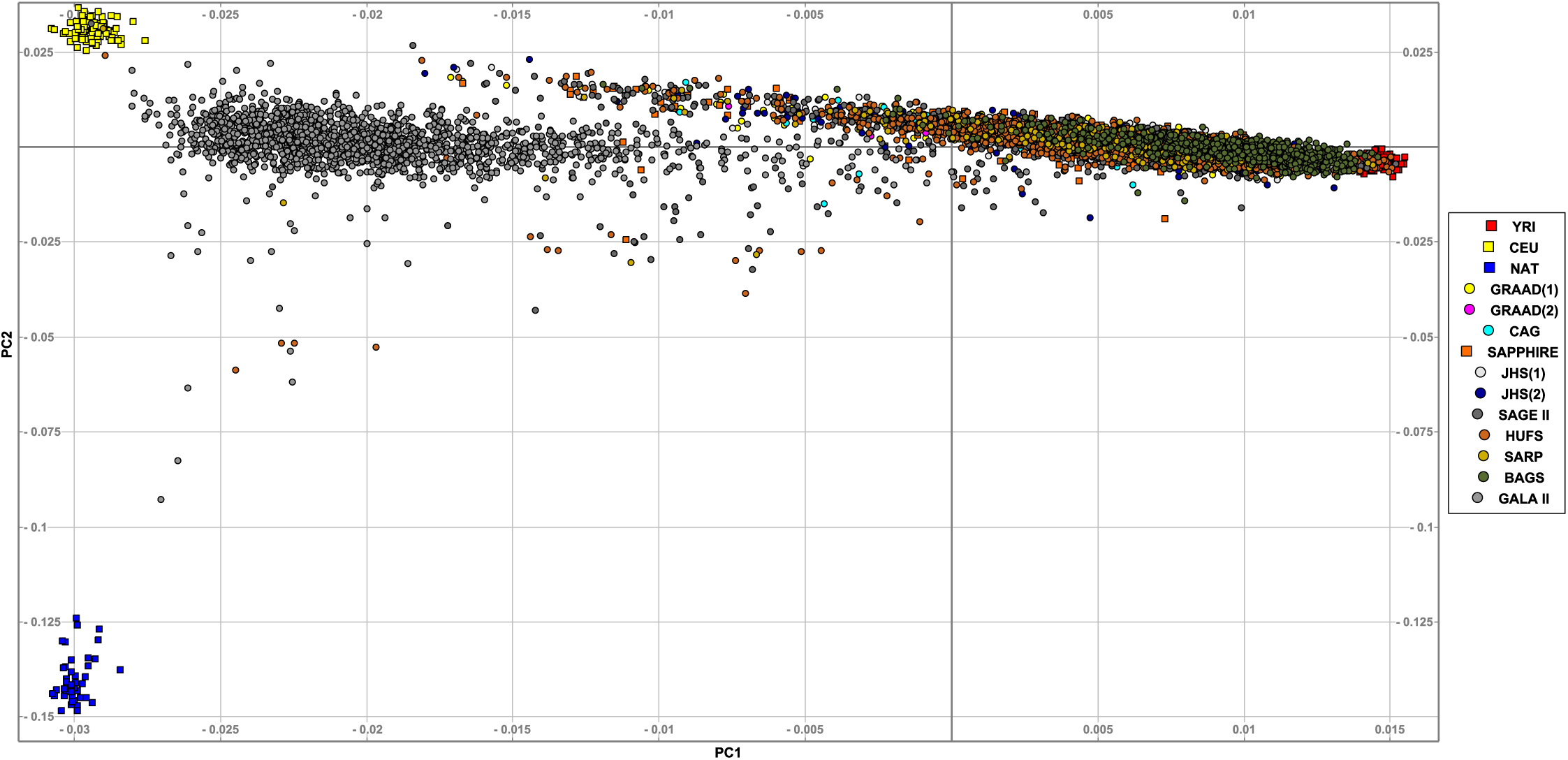
CAAPA principal components. This figure summarizes the first three principal components generated from a combined dataset of 16,410 overlapping SNPs in 84 African (YRI) and 84 European (CEU) TGP phase 3 subjects, 43 Native American (NAT) subjects and 9,170 CAAPA subjects. Three CAAPA outliers that clustered with CEU subjects (bottom leftmost corner of the plot) were identified based on this principal component analysis, and were excluded from the association analysis.

**Supplementary Figure S2:**
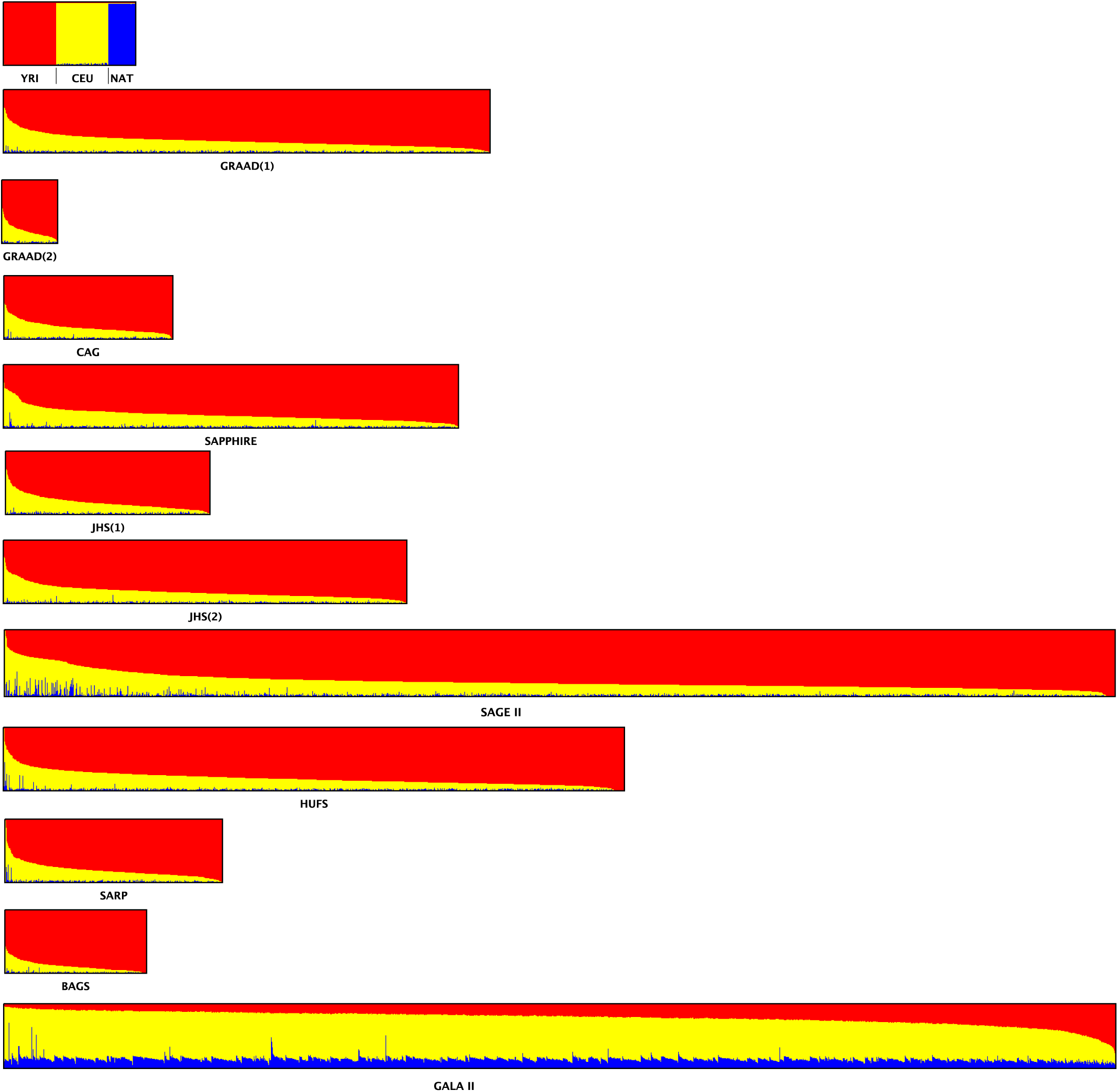
CAAPA ADMIXTURE estimates. This figure summarizes the genome-wide proportions of ancestry for K=3 populations, as estimated by the software program ADMIXTURE. A combined dataset of 16,410 overlapping SNPs in 84 African (YRI, red) and 84 CEU (European, yellow) TGP phase 3 subjects, 43 Native American (NAT, blue) subjects and 7,861 putatively unrelated CAAPA subjects were used to generate these estimates.

**Supplementary Figure S3:**
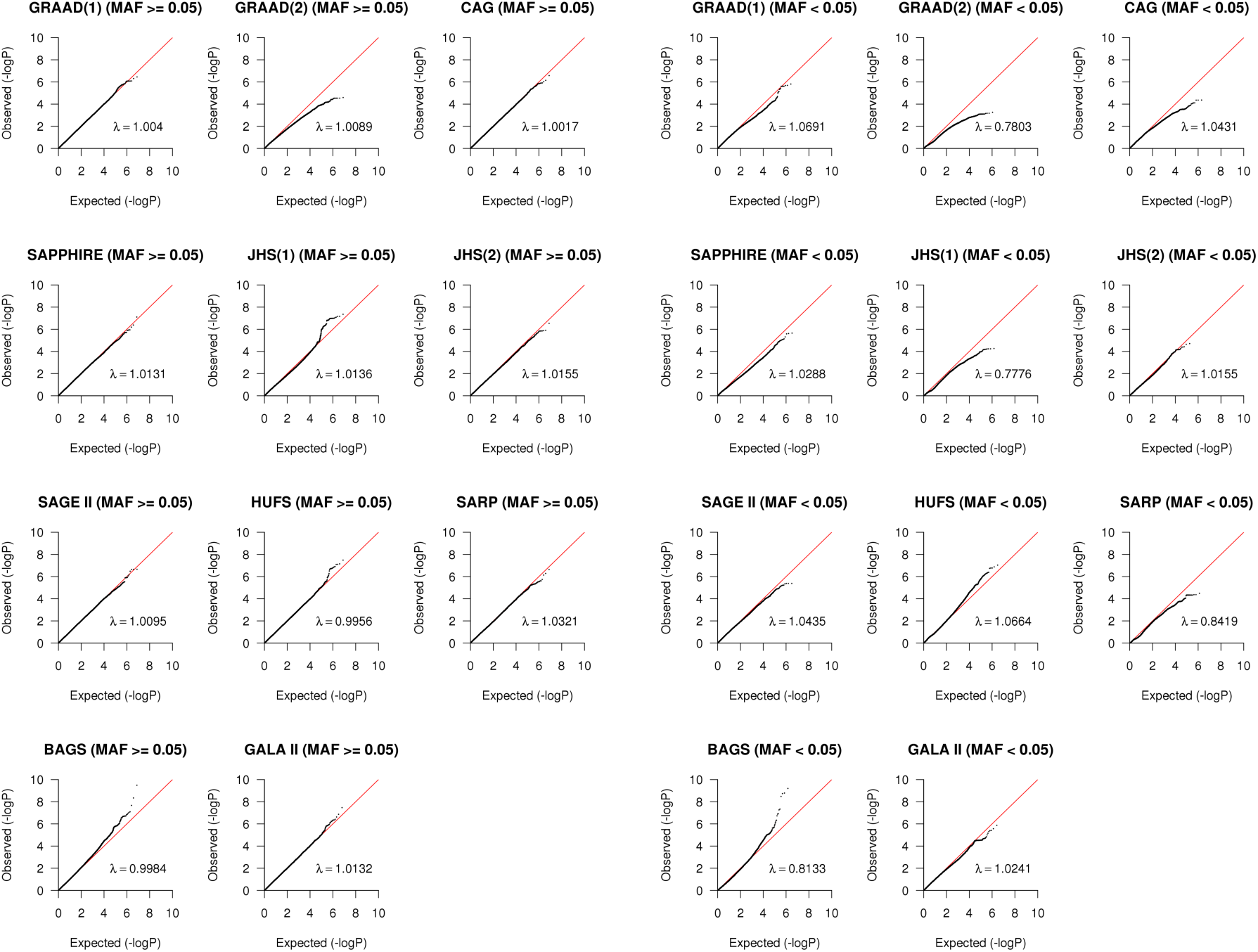
QQ plots summarizing the distribution of p-values per dataset, stratified by minor allele frequency (MAF). SNPs with minor allele count ≤ 6 were filtered out from the result sets. Inflation factors were calculated by transforming p-values to 1 degree of freedom (df) Chi-square statistics, and dividing the median of these statistics by the median of the theoretical Chi-square (1 df) distribution.

**Supplementary Figure S4:**
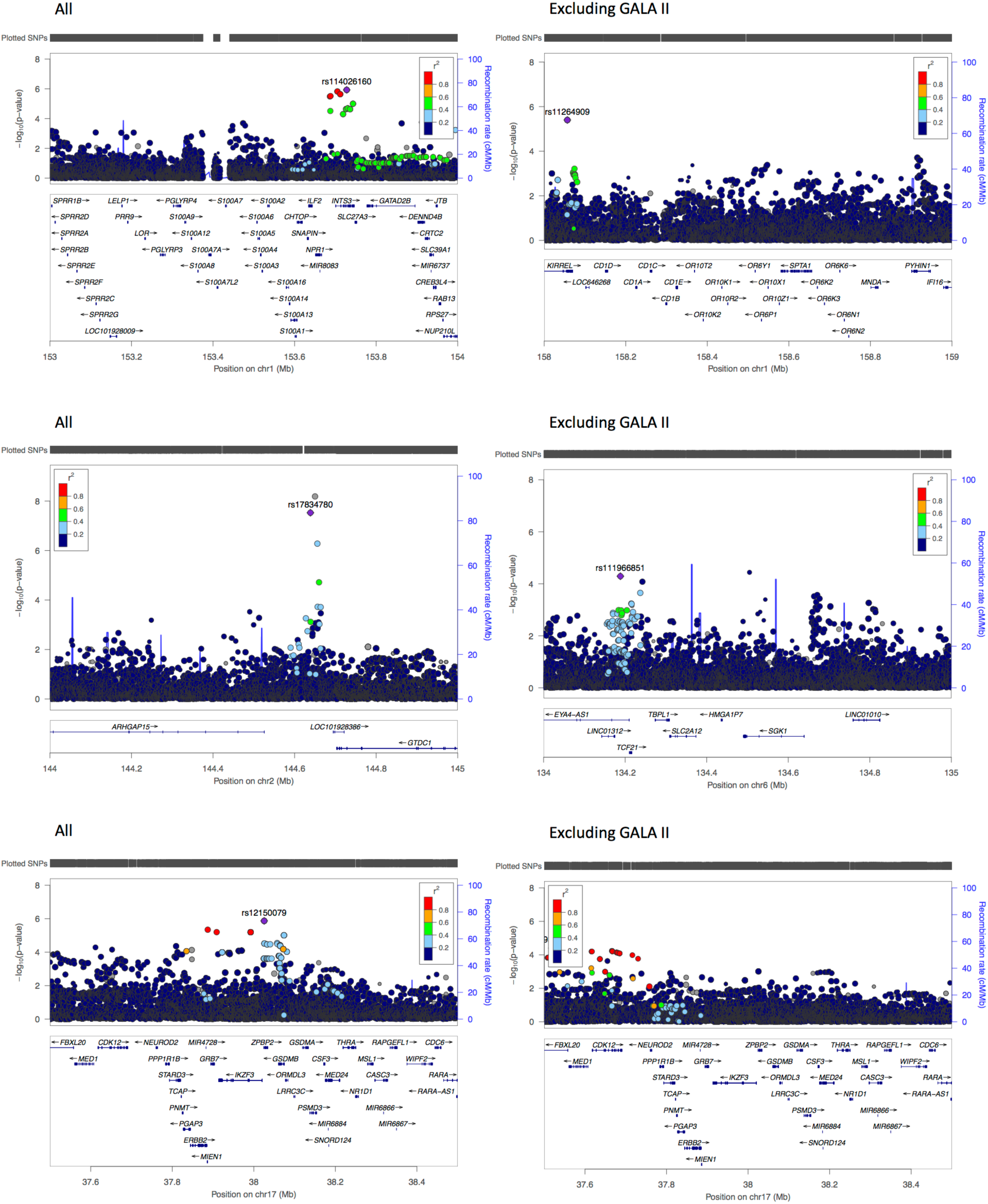
Locus zoom plots of the association p-values for the chromosome 1q21.3, 1q23.1, 2q22.3, 6q23.2 and 17q21 regions. In brackets, the meta-analysis result set used to generate the figures: “all” is the result set including all datasets, “excluding GALA II” is the result set excluding GALA II.

**Supplementary Figure S5:**
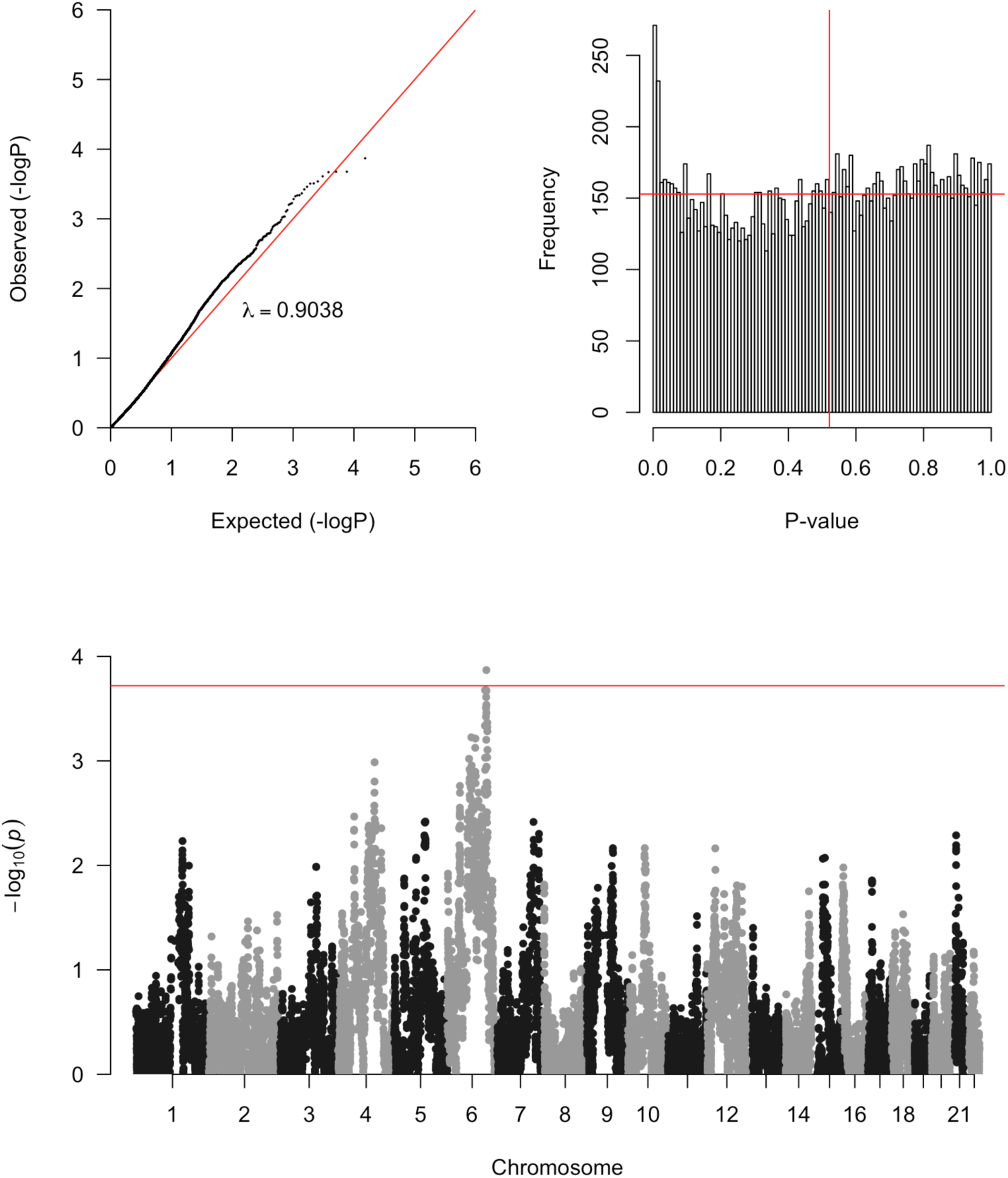
Summary of the distribution of admixture mapping p-values. A QQ plot is shown in the top panel, followed by a histogram of p-values, and the Manhattan plot is shown in the bottom panel. The inflation factor was calculated by transforming p-values to 1 degree of freedom (df) Chi-square statistics, and dividing the median of these statistics by the median of the theoretical Chi-square (1 df) distribution. The vertical red line in the histogram indicates the median p-value, and the horizontal red line represent the theoretical uniform distribution that the p-values are expected to follow. The horizontal red line in the Manhattan plot represents the p-value significance threshold of 1.9 × 10^−4^.

**Supplementary Figure S6:**
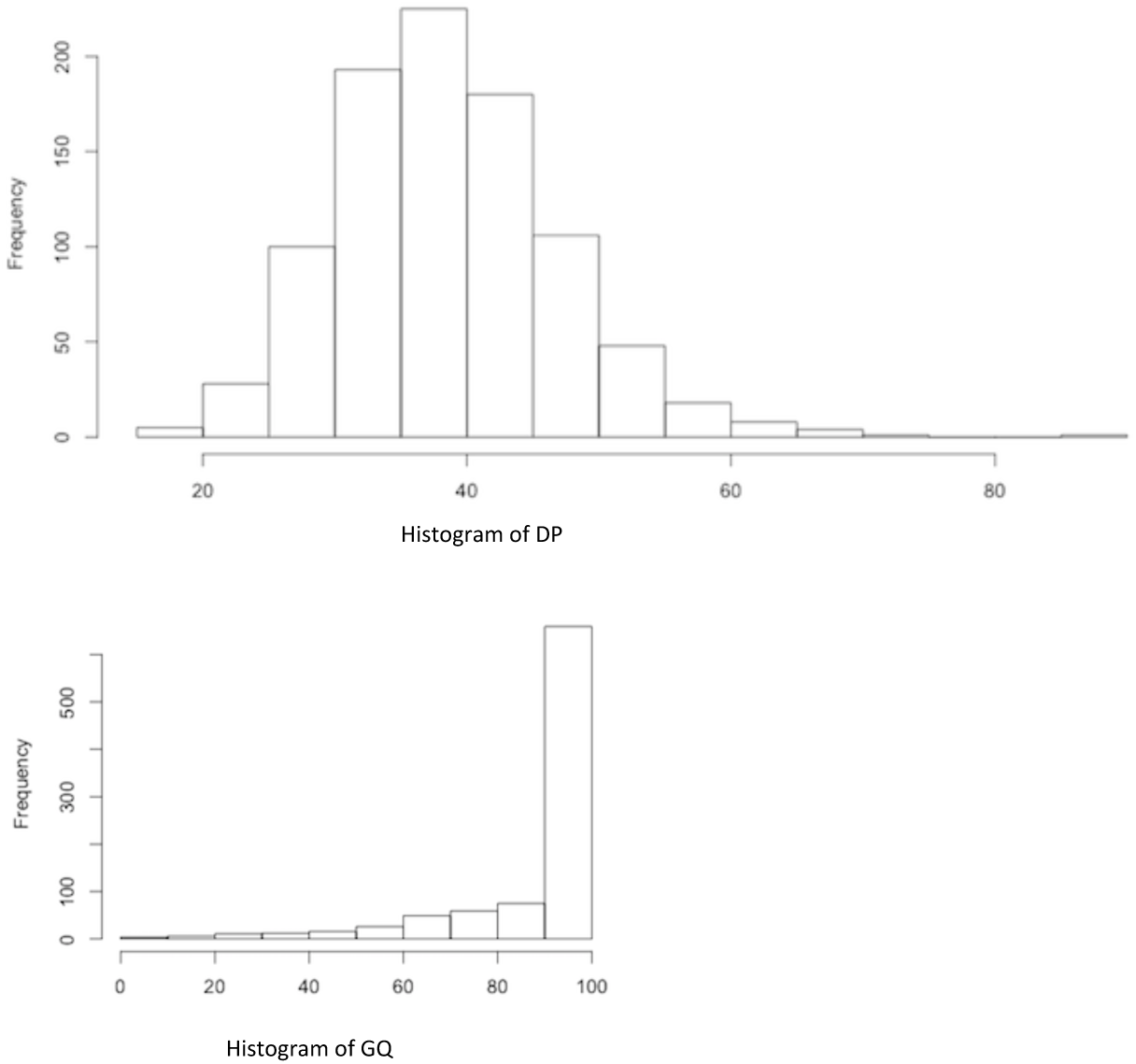
Summary of the distribution of the DP and GQ variant call quality metrics in the CAAPA WGS reference panel, for rs787160.

**Supplementary Figure S7:**
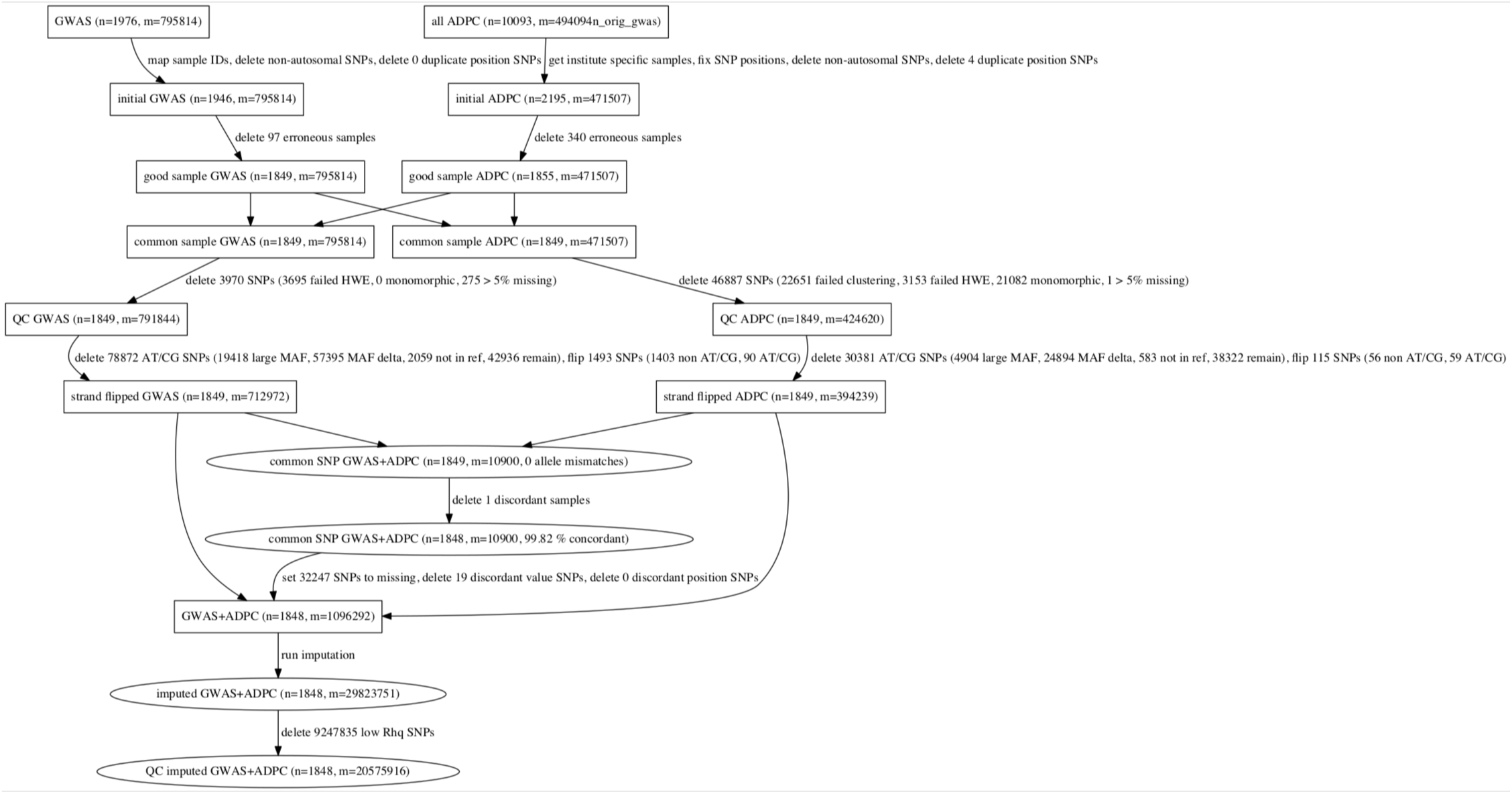
Flow diagram of GWAS+ADPC dataset merging and imputation. This figure depicts the steps used to merge and impute the GWAS backbone and ADPC datasets, as applied to HUFS.

**Supplementary Figure S8:**
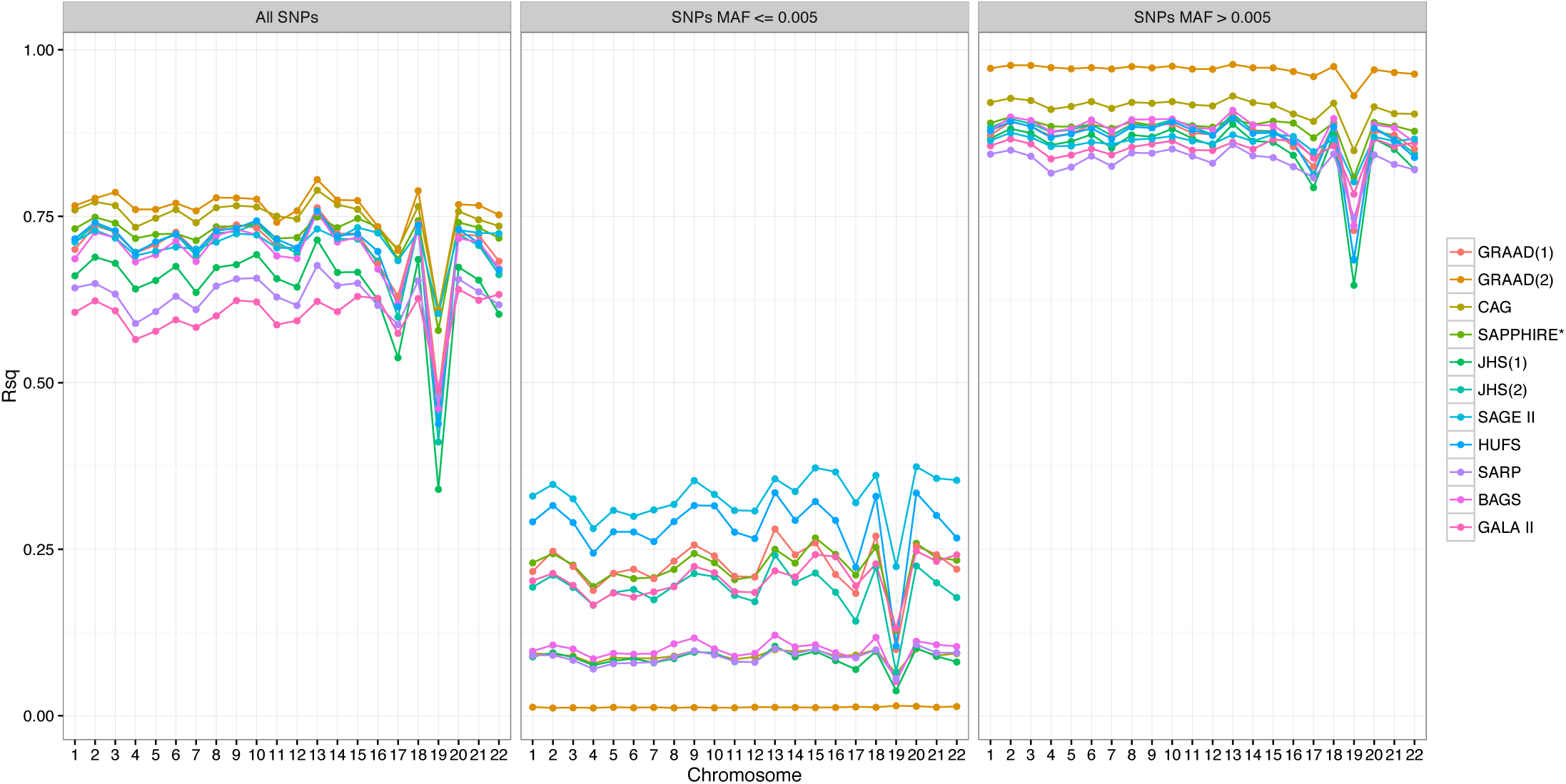
Median Rsq values by chromosome and dataset. The first panel represents the median Rsq values across all SNPs, the second panel represents the median Rsq values when sub-setting to SNPs with MAF ≤ 0.005, and the third panel represents the median Rsq values when sub-setting to SNPs with MAF > 0.005.

**Supplementary Figure S9:**
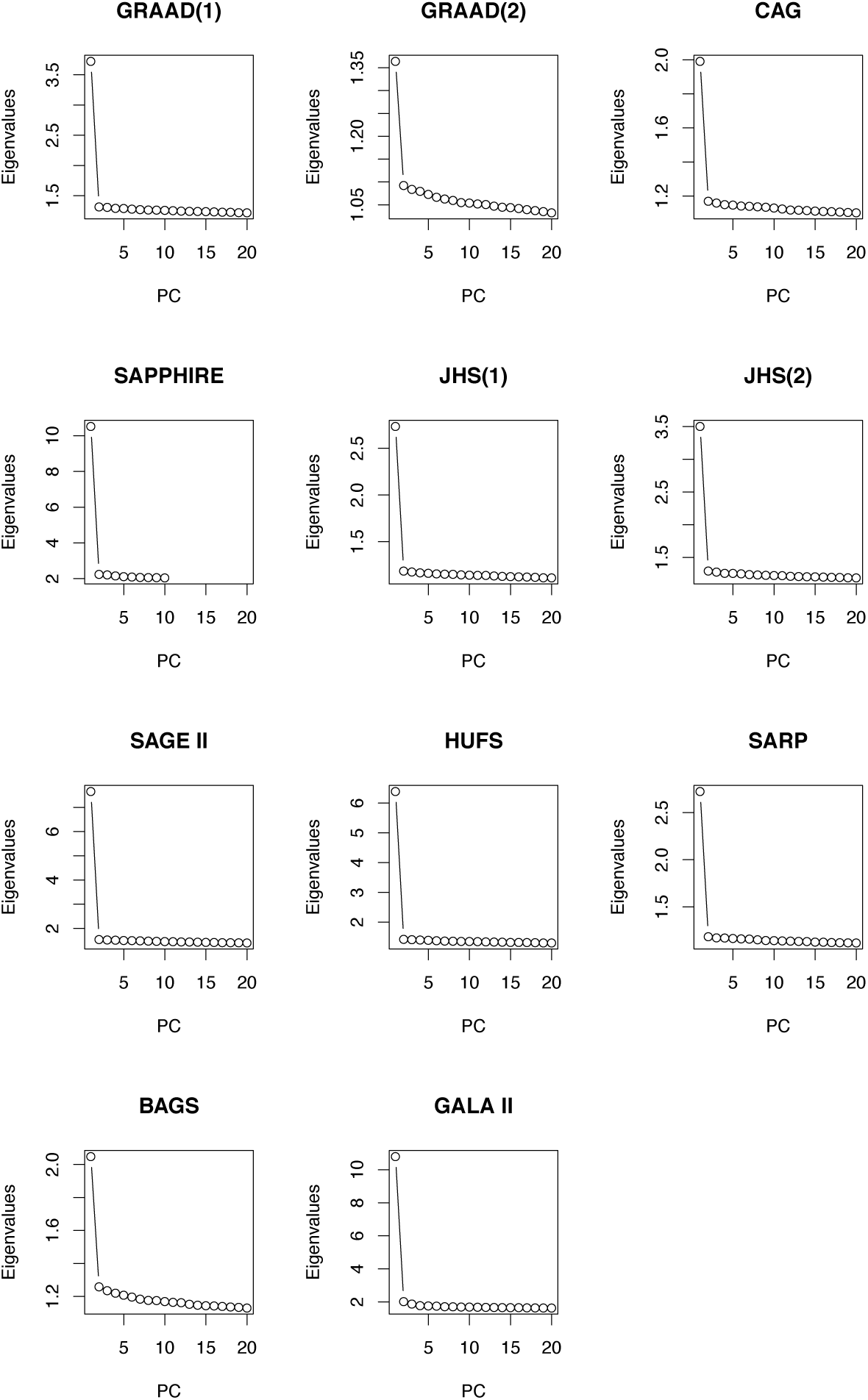
Scree plots of the within-dataset principal component analysis. Principal components were calculated using GENESIS separately for each dataset, excluding reference population panels. The plots show the eigenvalues of each principal component. Principal component 1 explains most of the variance in the data for all the CAAPA datasets.

**Supplementary Figure S10:**
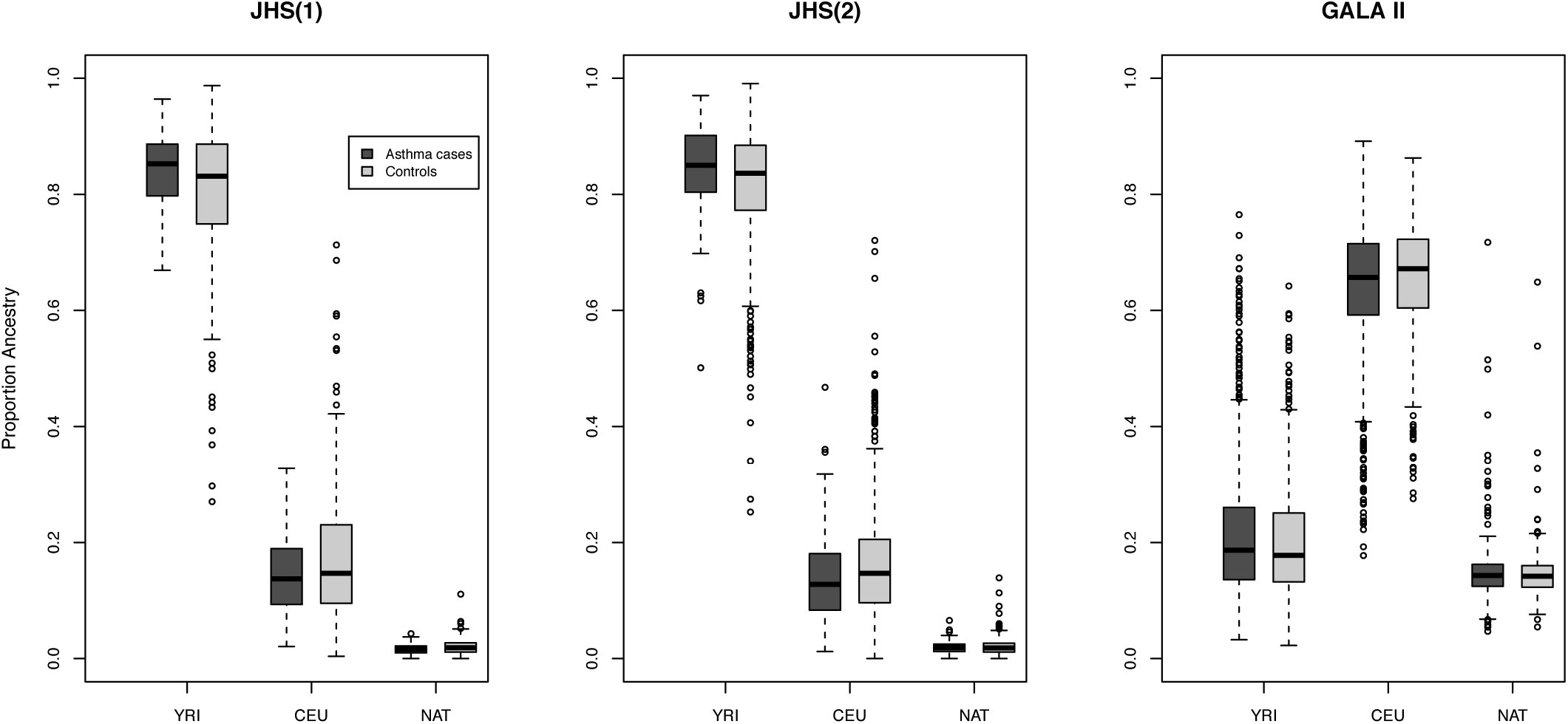
Distribution of ADMIXTURE estimates in the JHS and GALA II datasets. The association between ADMIXTURE estimates of genome-wide ancestry and case-control status was tested using logistic regression, separately for each dataset and ancestry. The ADMIXTURE estimates shown in Supplementary Figure S4 were used to generate these boxplots and association p-values.

**Supplementary Figure S11:**
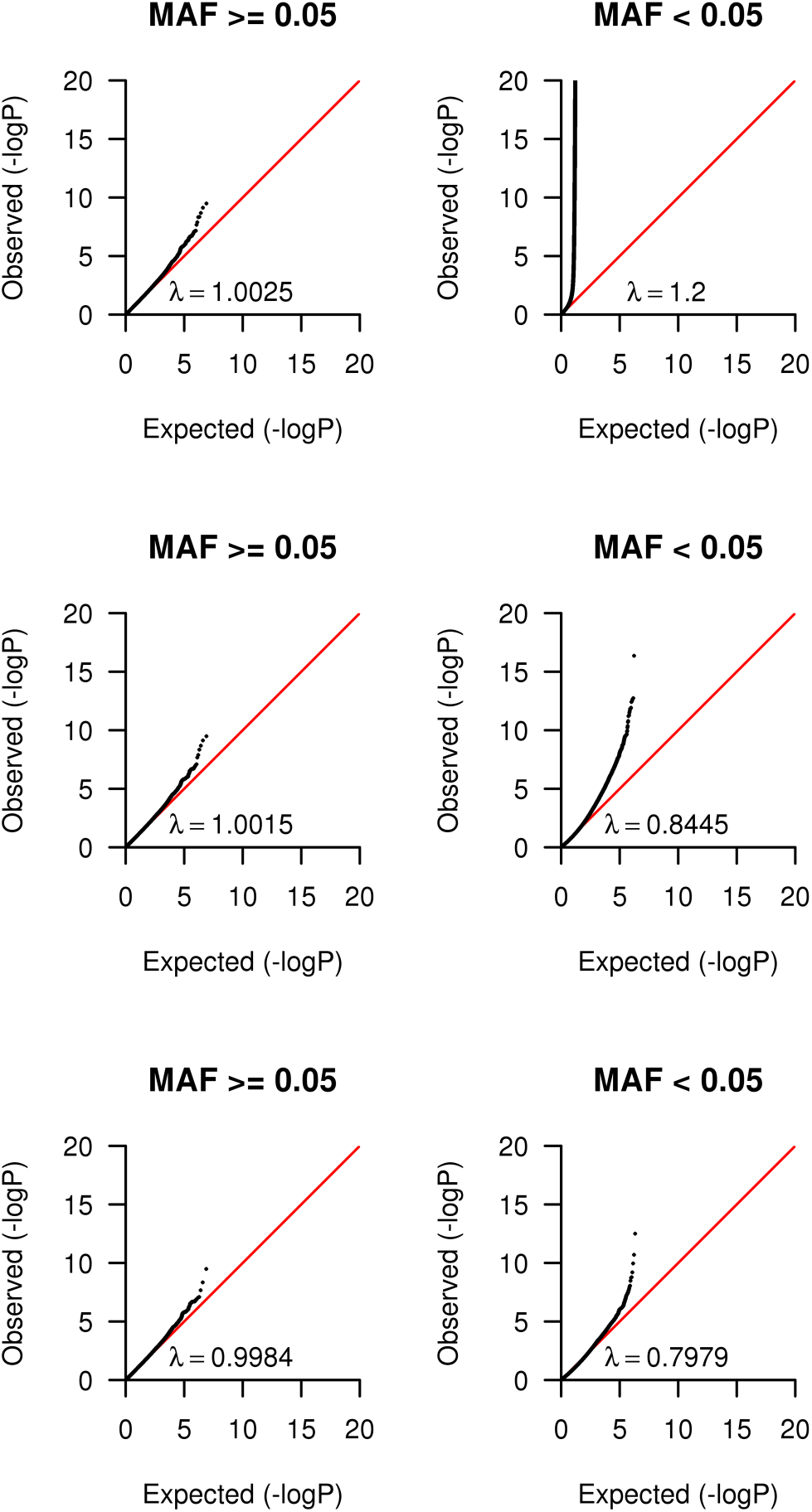
Summary of the distribution of BAGS p-values, estimated using MQLS, for different Rsq filters. Inflation factors were calculated by transforming p-values to 1 degree of freedom (df) Chi-square statistics, and dividing the median of these statistics by the median of the theoretical Chi-square (1 df) distribution. A.) QQ plots filtered using Minimac Rsq and MAF estimates. SNPs with MAF ≤ 0.005 were deleted if Rsq ≤ 0.5 and SNPs with MAF > 0.005 were deleted if Rsq ≤ 0.3. B.) QQ plots filtered using MQLS Rsq and control MAF estimates, that takes into account the correlation between subjects due to relatedness. SNPs with MAF ≤ 0.005 were deleted if Rsq ≤ 0.5 and SNPs with MAF > 0.005 were deleted if Rsq ≤ 0.3. C.) QQ plots filtered using MQLS Rsq and control MAF estimates. SNPs with MAF ≤ 0.005 were deleted if Rsq ≤ 0.7 and SNPs with MAF > 0.005 were deleted if Rsq ≤ 0.5. This filter was applied prior to including BAGS in the meta-analysis.

**Supplementary Figure S12:**
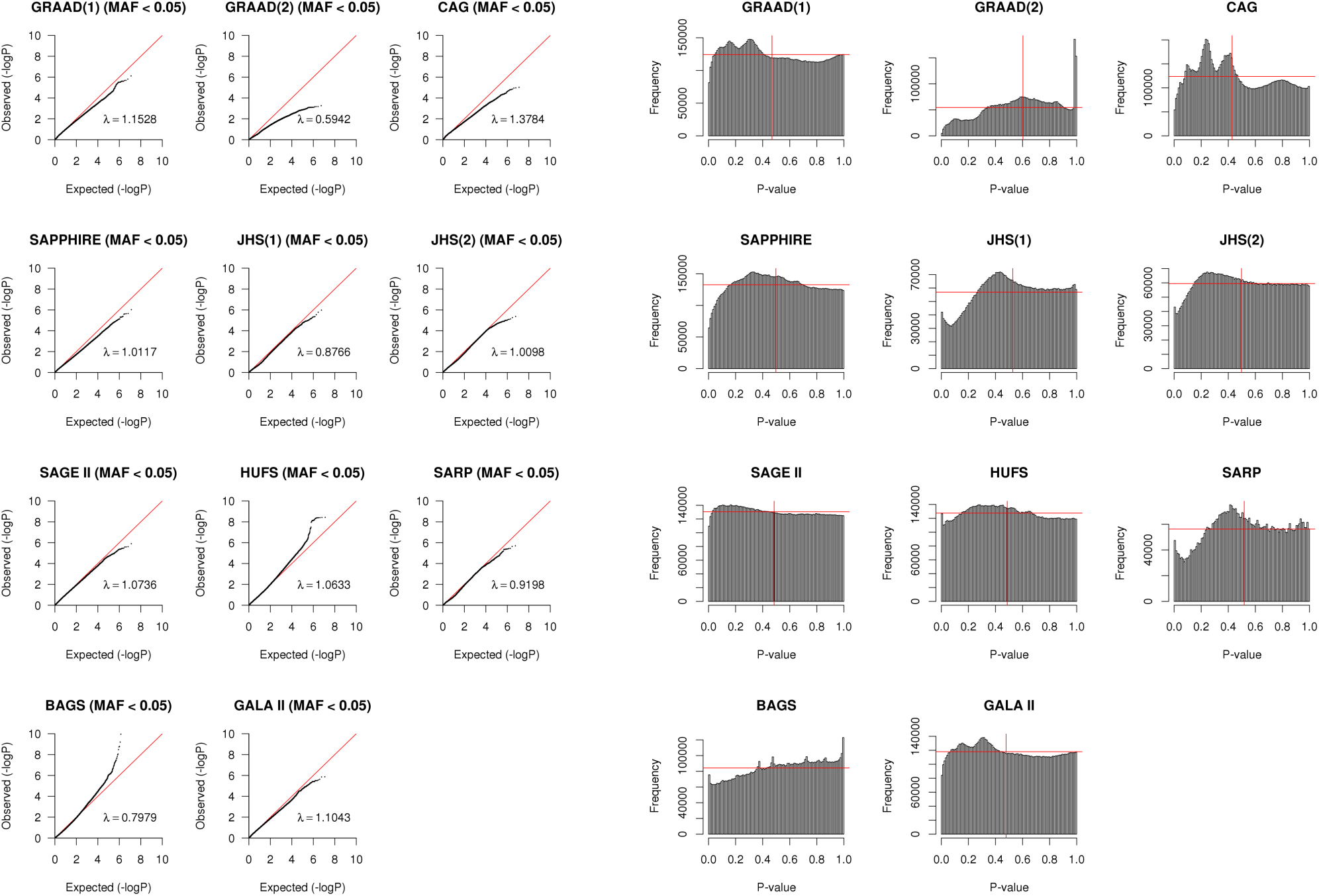
Summary of the distribution of p-values for SNPs with MAF < 0.05, prior to applying a minor allele count (MAC) filter. QQ plots and histograms of p-values are shown in the left and right panels respectively. Inflation factors were calculated by transforming p-values to 1 degree of freedom (df) Chi-square statistics, and dividing the median of these statistics by the median of the theoretical Chi-square (1 df) distribution. The vertical red lines in the histograms indicate the median p-value, and the horizontal red lines represent the theoretical uniform distribution that the p-values are expected to follow.

**Supplementary Figure S13:**
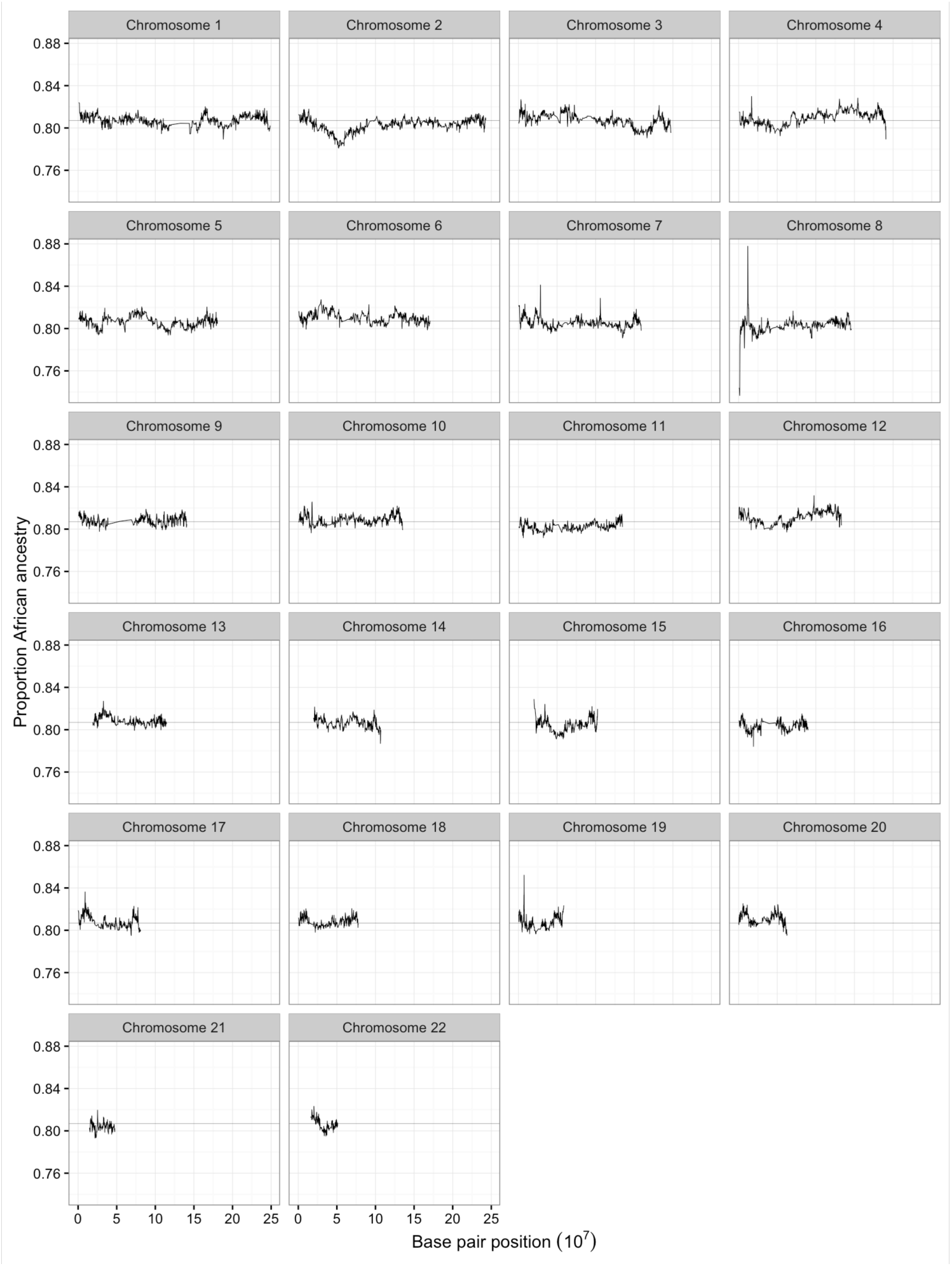
The mean proportion of Africa ancestry across the genome of 7, 146 CAAPA subjects, for the 15,824 local ancestry segments inferred by RFMix. The mean local African ancestry for a segment is the proportion of haplotypes called as African for that segment.

**Supplementary Figure S14:**
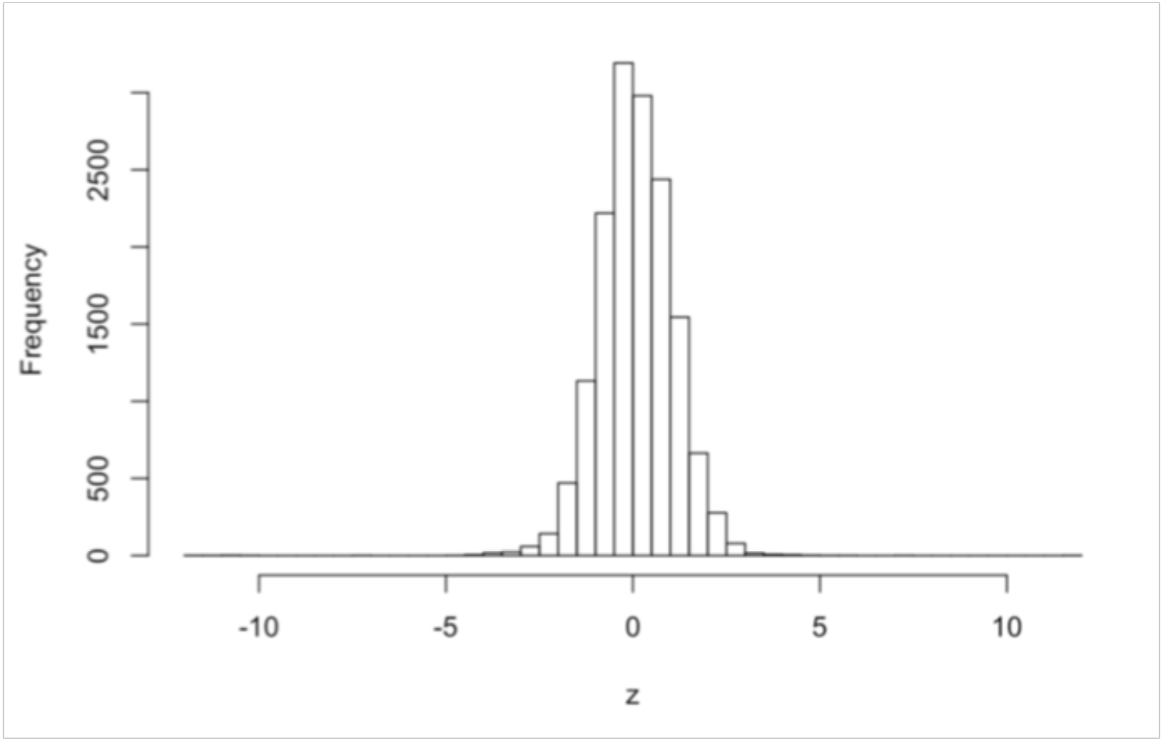
Histogram of the standardized deviations between mean local African ancestry and mean genomewide African ancestry, for the 15,824 local ancestry segments inferred by RFMix. The mean local African ancestry for a segment is the proportion of haplotypes called as African for that segment. The mean genome-wide African ancestry was calculated by dividing the number of SNPs called by RFMix as African, by the total number of SNPs for which local ancestry was called, across all genomes.

**Supplementary Figure S15:**
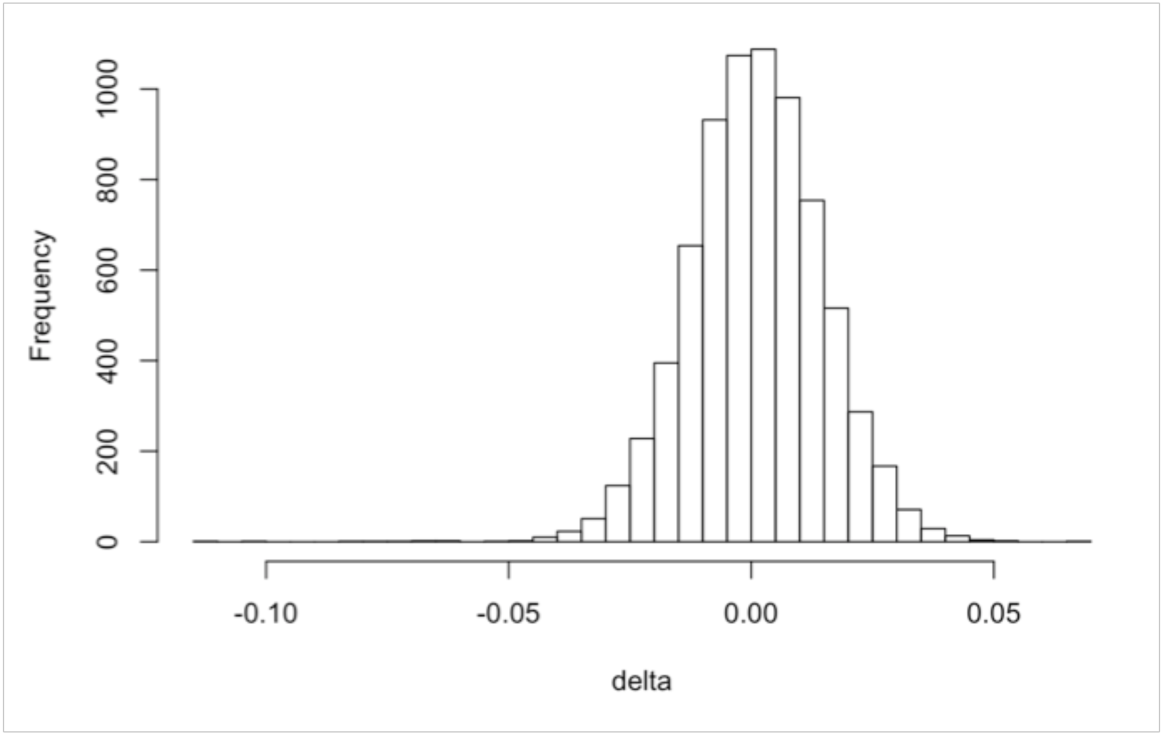
Histogram of the differences between genome-wide ancestry estimated using ADMIXTURE, and genome-wide ancestry estimated using RFMix. The difference (delta) was calculated for each of the 7,416 individuals for which local ancestry was called. The per-individual RFMix mean genome-wide African ancestry was calculated by dividing the number of SNPs called by RFMix as African, by the total number of SNPs for which local ancestry was called across the genome, for that individual.

### SUPPLEMENTAL TABLES

**Supplementary Table S1:**
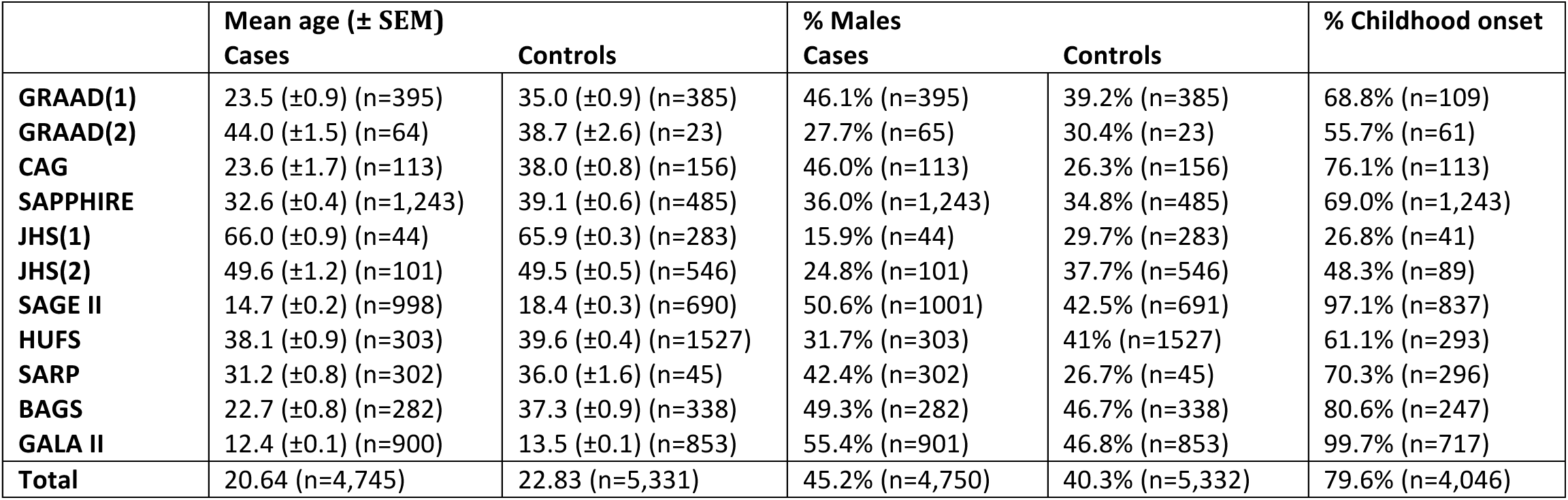
Clinical characteristics summary. The distribution of age, sex and age of asthma onset is summarized in this table by dataset used in the meta-analysis. SEM=standard error of the mean. Childhood onset asthma is defined as having been diagnosed with asthma before 16 years of age. The information summarized in this table was not available for all subjects included in the association analysis; therefore the number of subjects (n) used to calculate these statistics is given in brackets.

**Supplementary Table S2:**
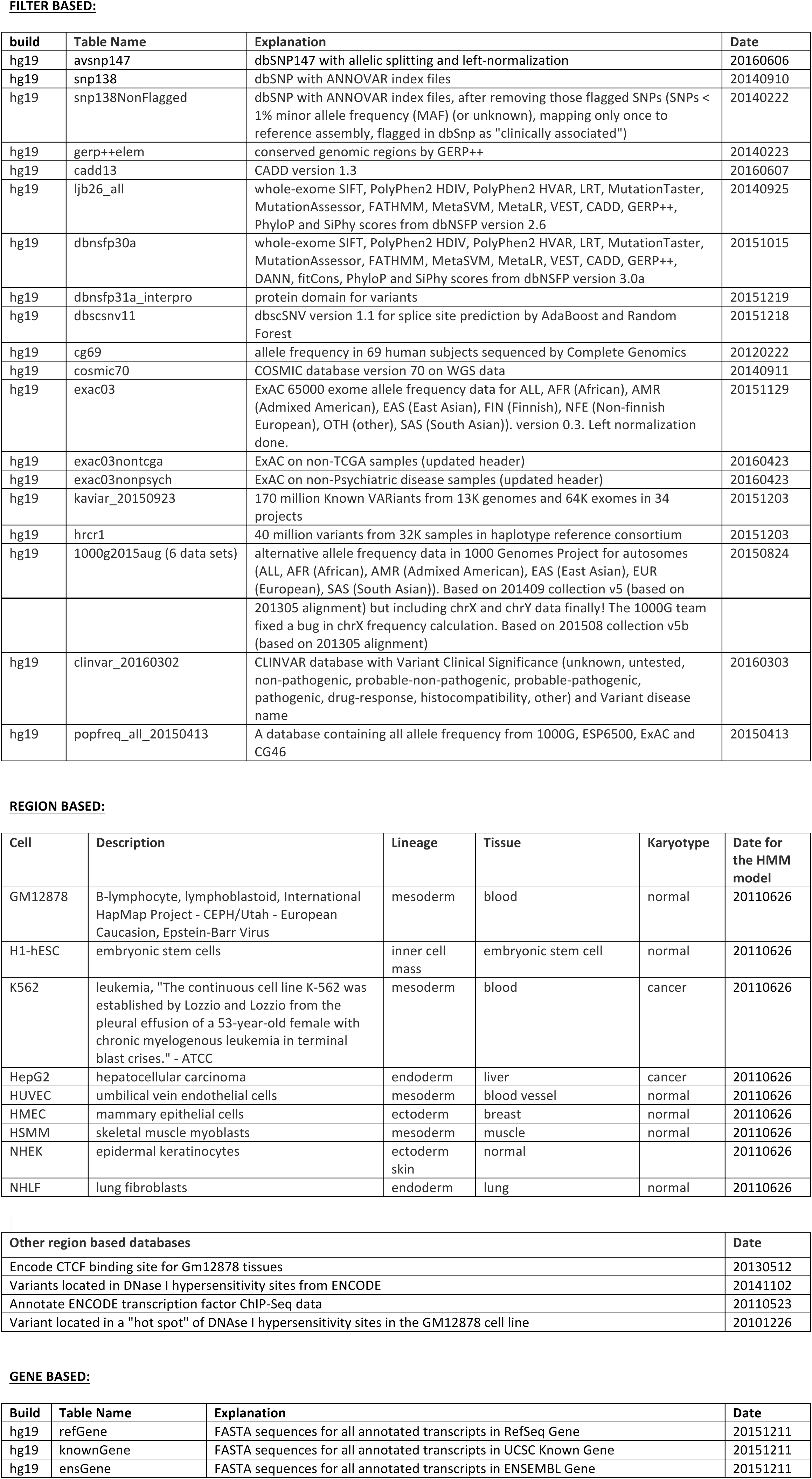
ANNOVAR file versions.

**Supplementary Table S3:**
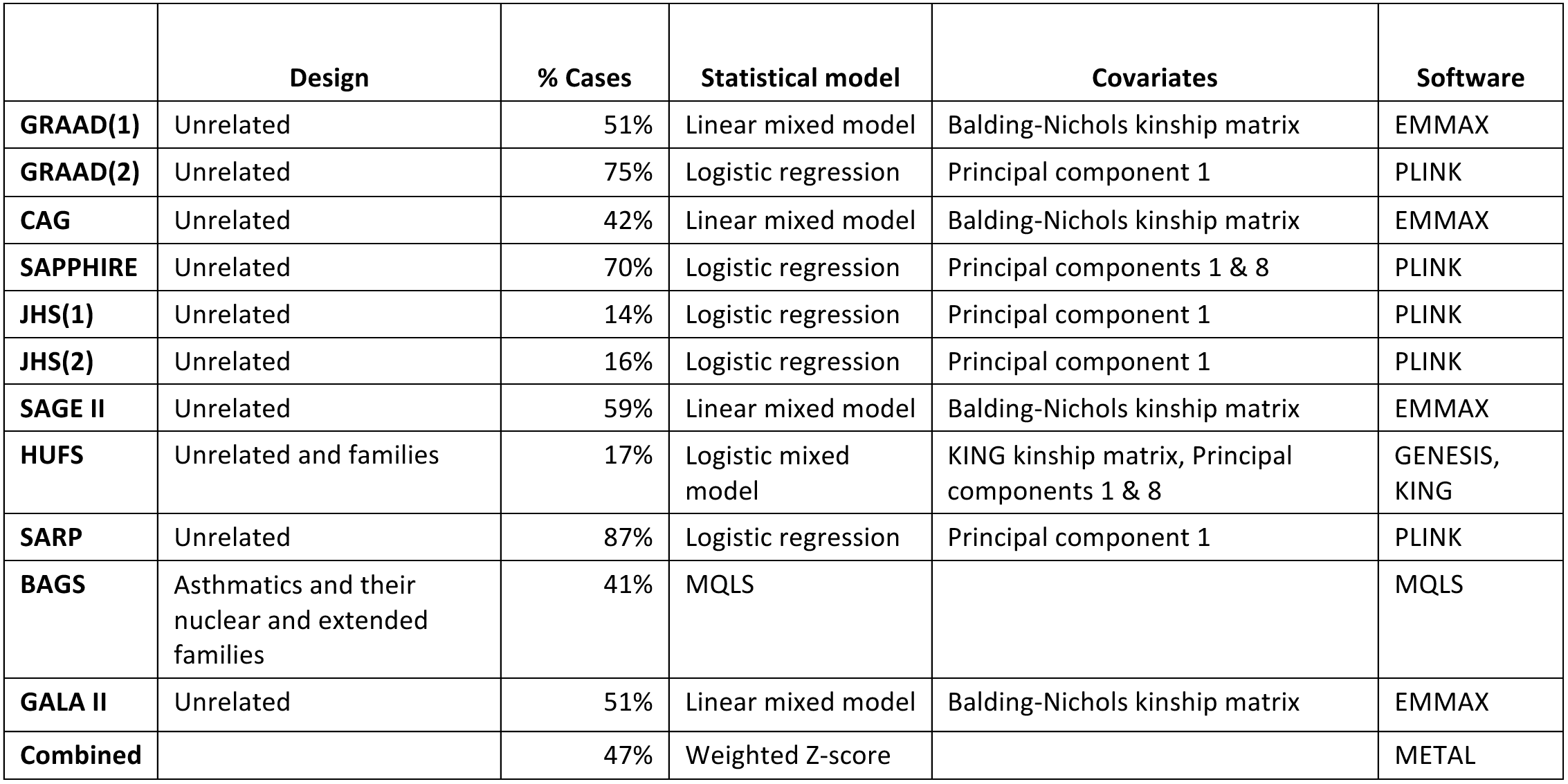
Statistical models used for association testing. For the logistic regression models that included only fixed effects (no kinship matrix), for each pair of subjects that shared a kinship coefficient > 0.25, the subject with the least represented phenotype were deleted. For all logistic regression models, the first principal component as well as any of the top 10 principal components that were associated with asthma status (p-value < 0.05), were included as covariates.

**Supplementary Table S4:**
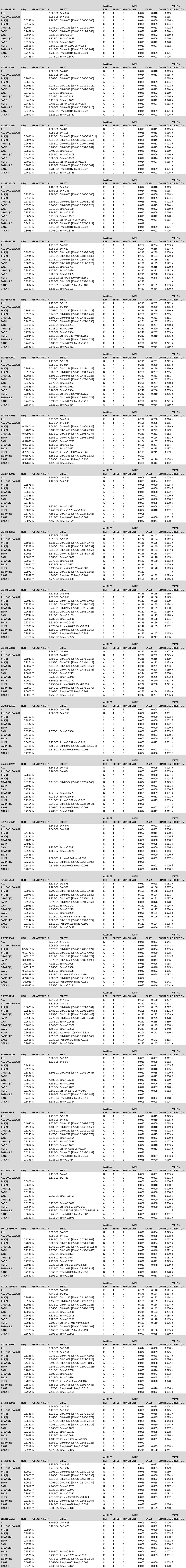
Summary of association results by meta-analysis result set and individual CAAPA dataset. The ALL rows refers to the meta-analysis result including all CAAPA datasets. The ALL EXCL GALA II rows refer to the meta-analysis result excluding the GALA II dataset. SNPs are identifed by their chr:hg19_position. SNPs with the following criteria are summarized: SNPs with p-values < 10-6 in either of the meta-analysis result sets that include or exclude GALA II, and SNPs that fall within an asthma candidate gene region and that pass a multiple testing significance threshold for that gene region in either of the two meta-analysis result sets. Two SNPs close to the PYHIN1 gene, reported by the EVE meta-analysis, are also summarized (1:158932555 and 1:158932907 respectively).

**Supplementary Table S5:**
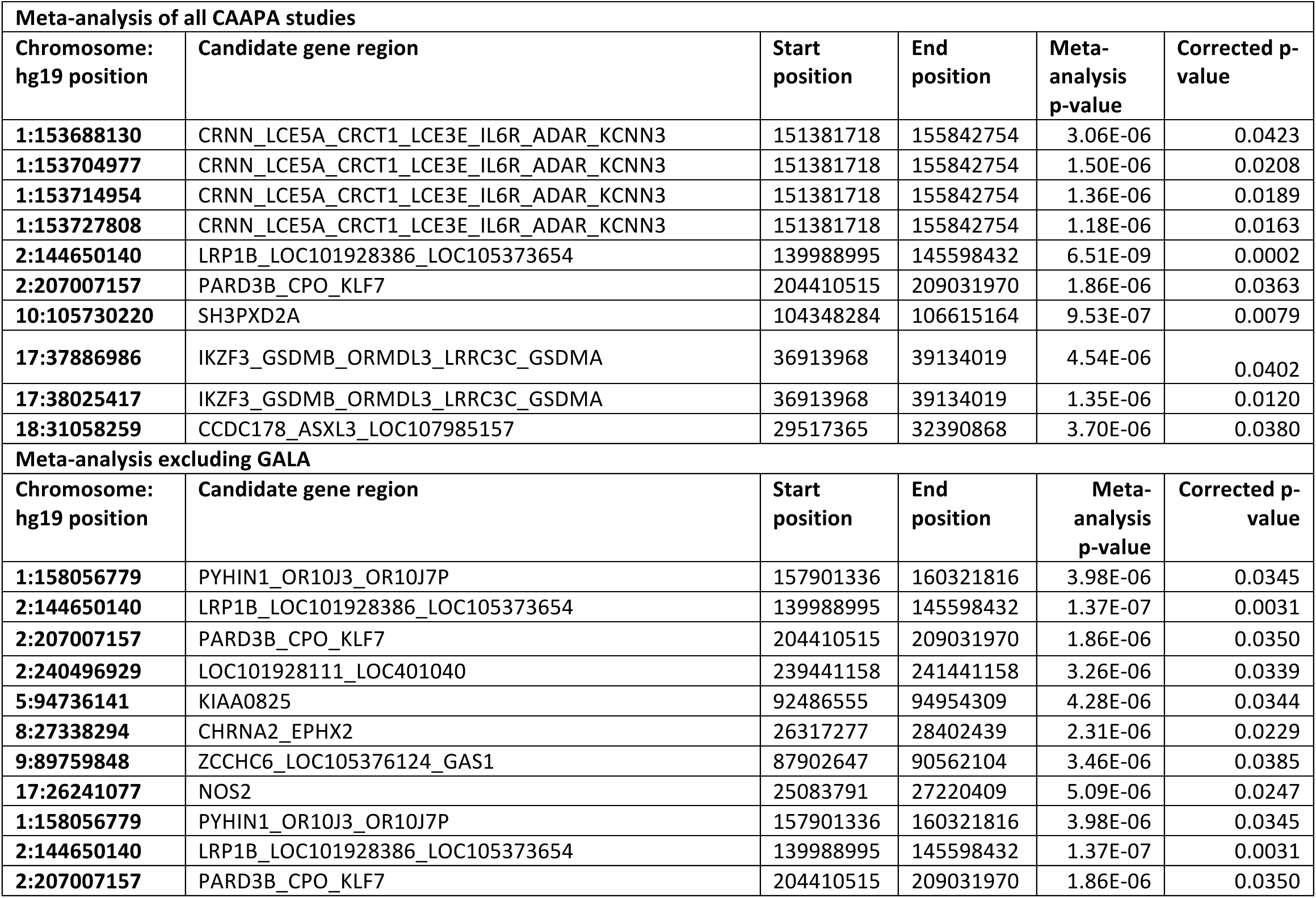
Candidate gene association results. This table summarizes the candidate genes that are associated with asthma after applying a Bonferroni correction using the number of independent markers in the candidate gene region. Results are shown for the meta-analysis of all the CAAPA studies, and the meta-analysis that excludes the GALA II study, due to the relatively higher European ancestry within GALA II.

**Supplementary Table S6:**
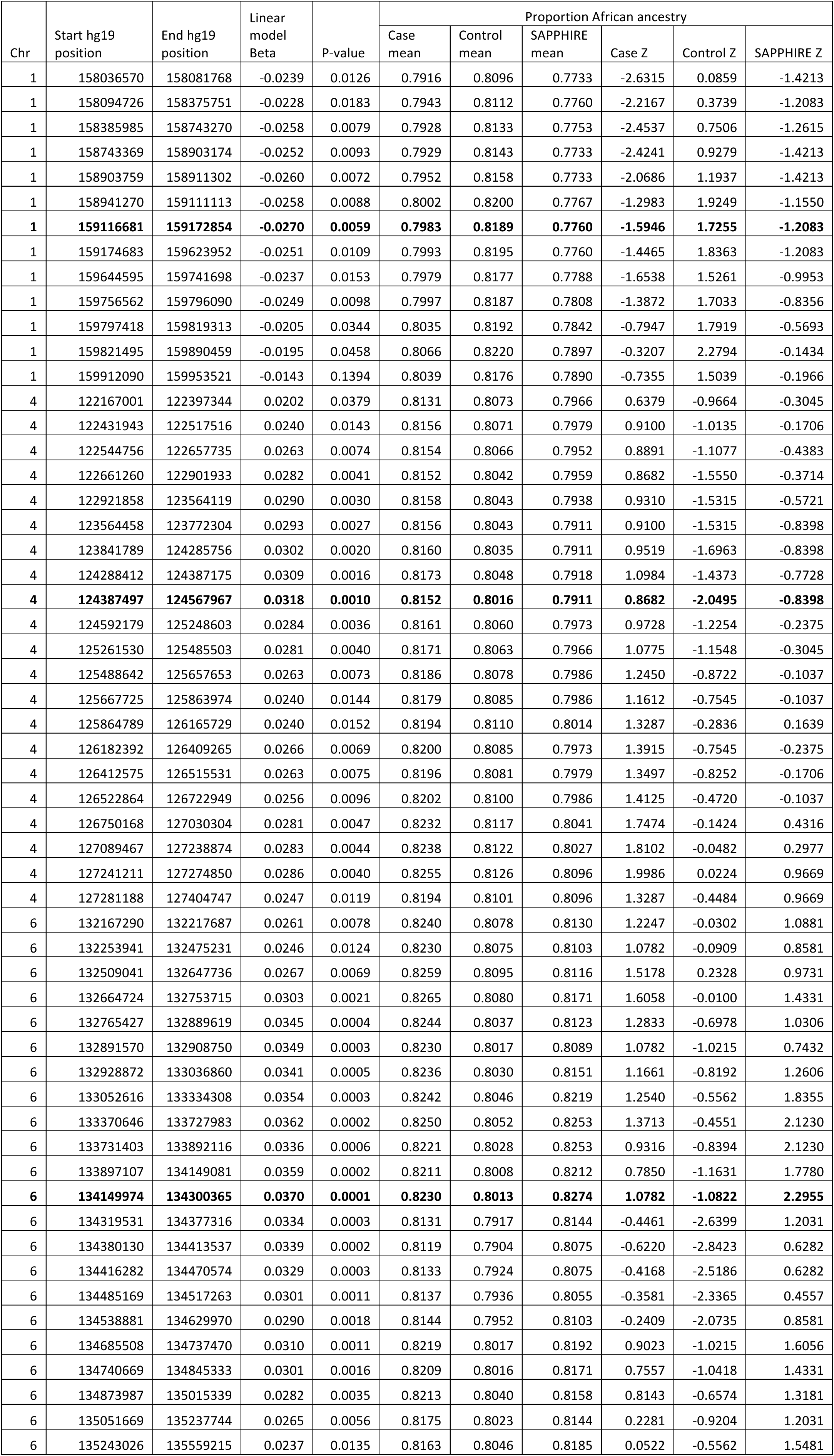
Summary of the admixture mapping peaks on chromosomes 1, 4 and 6. A linear mixed model (EMMAX software) was used to test for association between the number of copies of African ancestry (“dosage” value of 0, 1, 2) and case-control status (2,608 cases and 3,994 controls). Since ADPC data was only available for SAPPHIRE cases, the study was not included in the model. The most significant segments are highlighted in **bold font**. Z is the standardized deviation between the mean African ancestry of a local ancestry segment in cases/controls/SAPPHIRE, and the grand African ancestry mean in cases/controls/SAPPHIRE (the means and Z were calculated separately for each of these 3 groups).

**Supplementary Table S7:**
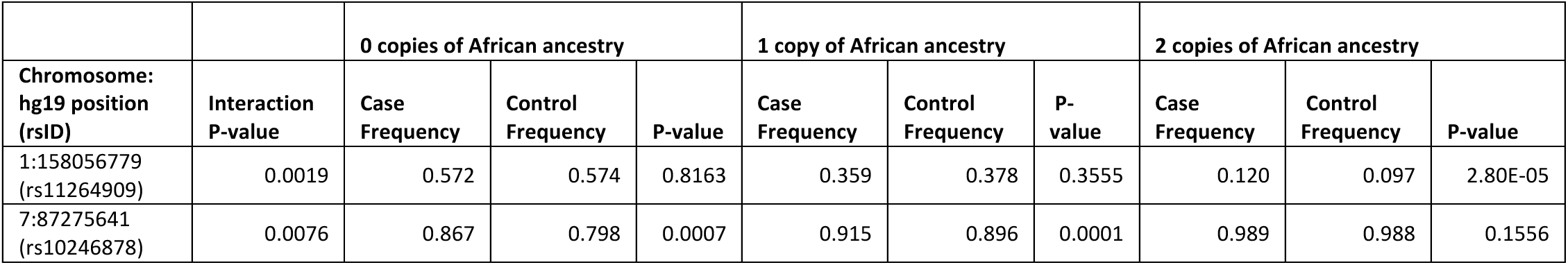
Summary of association tests stratified by local ancestry. A logistic regression interaction model was used to determine if an association differs by local ancestry. SNPs that had p-values < 10^−5^ in either of the meta-analyses (including vs. excluding GALA), or that were identified by our candidate gene analysis, were tested using this model. SNPs with MAF < 0.01 and with less than 8 studies contributing to the meta-analysis p-value were excluded from the test. Models with interaction p-values < 0.05 were further examined by stratifying the data set according to 0, 1 or 2 copies of African ancestry at that locus, and then using an additive allelic test for association within each strata, to test for differences in case and control allele frequency.

**Supplementary Table S8:**
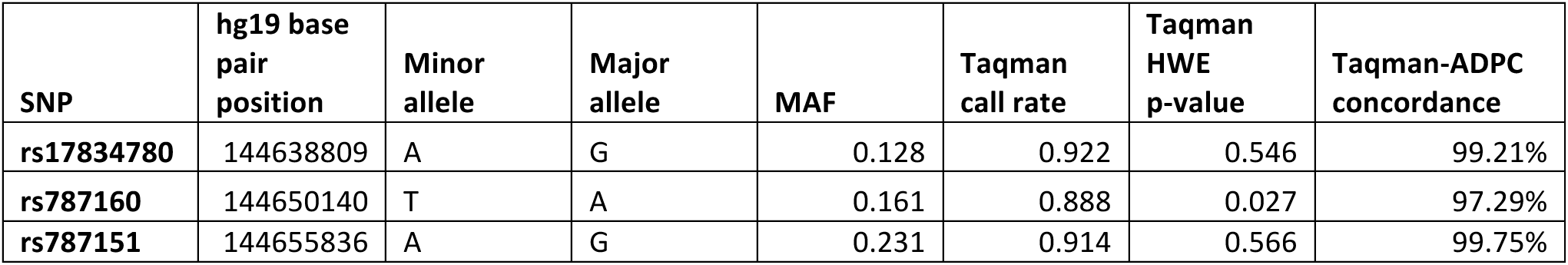
Summary of concordance between chromosome 2q22.3 Taqman and imputed genotypes. Samples from HUFS were typed using a TaqMan genotyping assay in order to do this comparison. The data is 98.76% concordant across 1792 patients with at least one SNP to compare.

**Supplementary Table S9:**
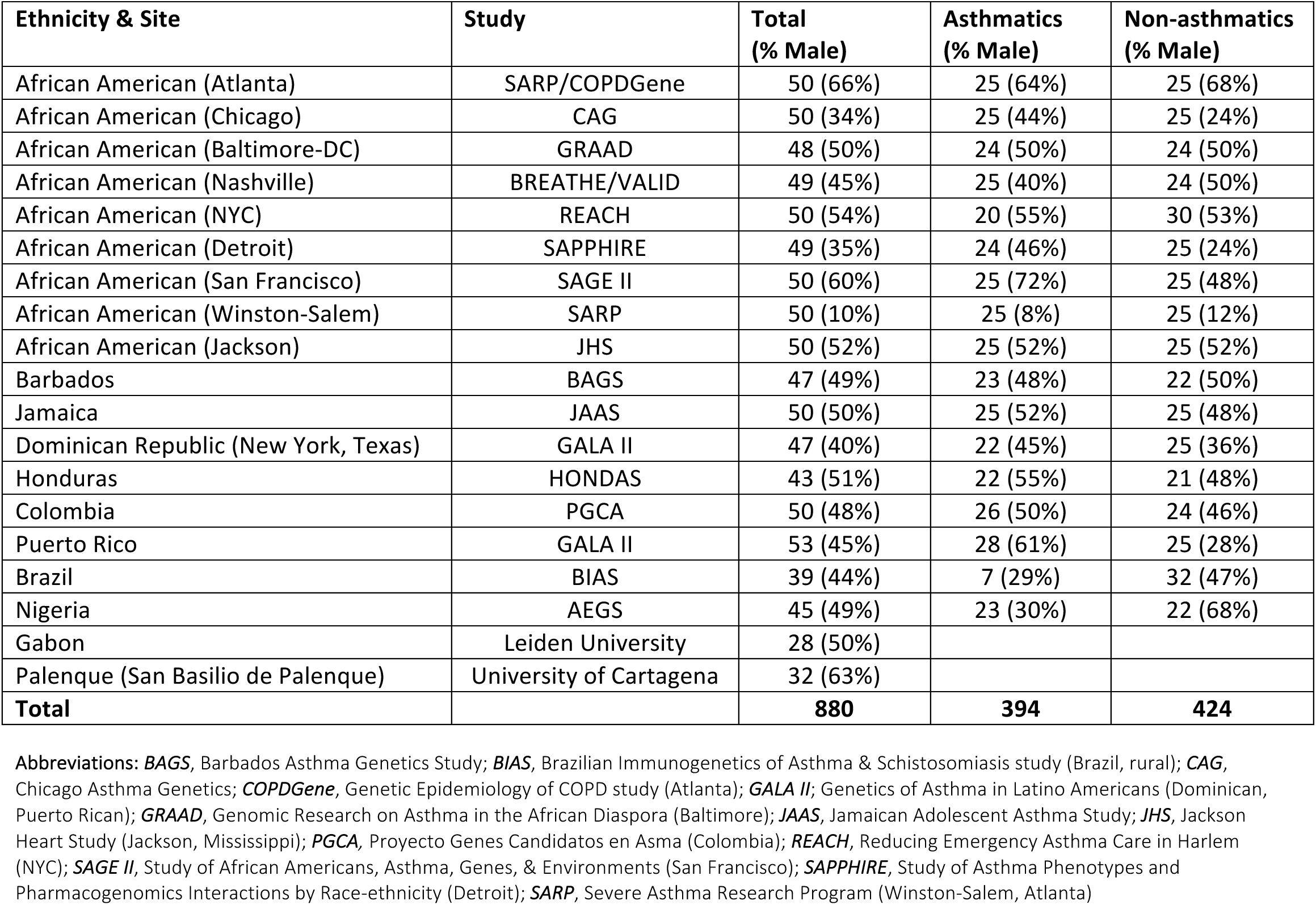
CAAPA whole genome sequence (WGS) reference panel. This table summarizes the ethnicity and recruitment sites of subjects used to build the CAAPA WGS reference panel.

**Supplementary Table S10:**
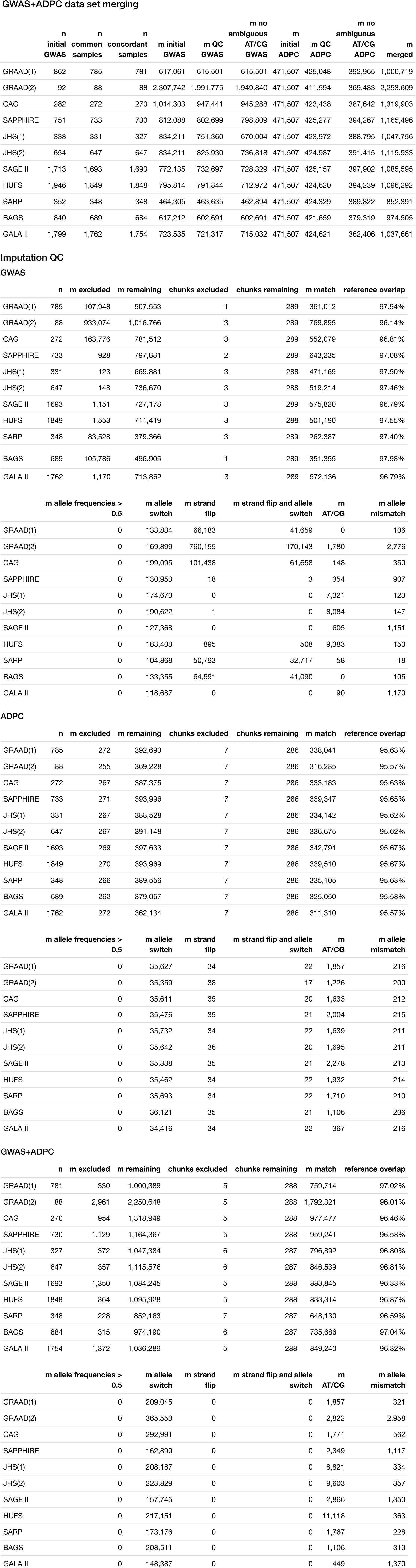
Summaries of GWAS+ADPC data set merging and imputation process results. The GWAS+ADPC data set merging table summarizes the number of subjects (n) and number of SNPs (m) available at different stages of the data merging process. The Imputation QC tables summarize the imputation QC report downloaded from the Michigan imputation server, for pre-imputation QC (Quality Control Only option) of the GWAS data sets and ADPC data sets, and impuation QC of the merged GWAS+ADPC data set.

**Supplementary Table S11:**
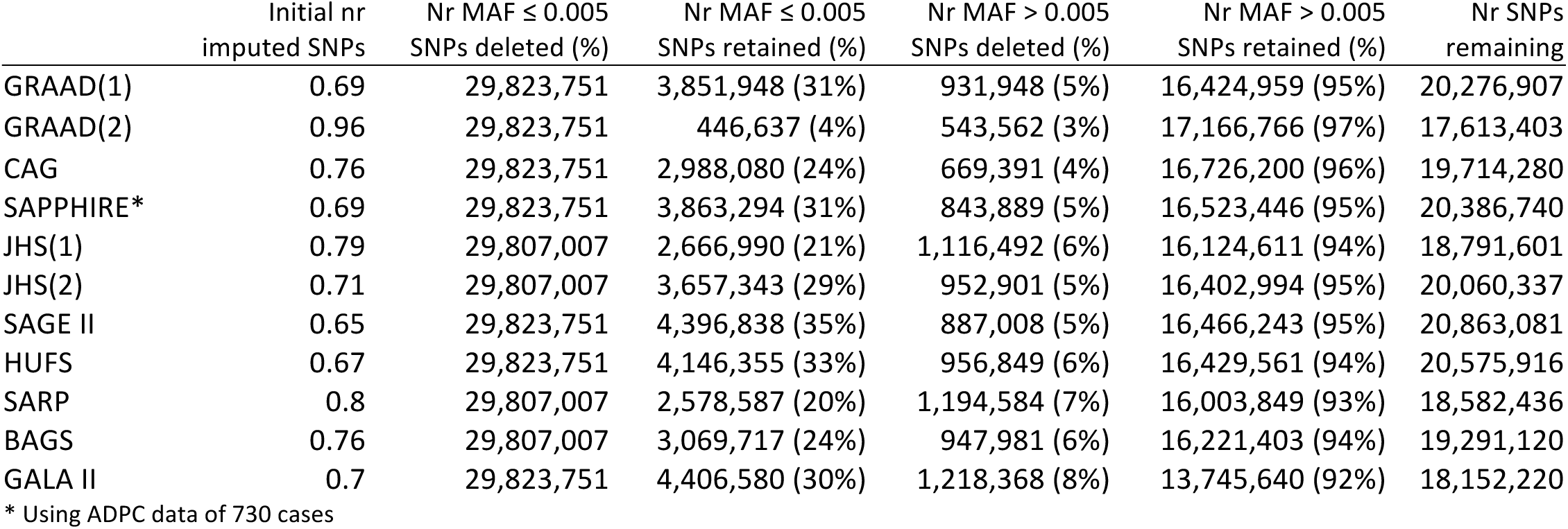
The proportion of SNPs deleted and retained in the imputed datasets by dataset and MAF category. SNPs with MAF ≤ 0.005 were deleted if Rsq ≤ 0.5 and SNPs with MAF > 0.005 were deleted if Rsq ≤ 0.3

**Supplementary Table S12:**
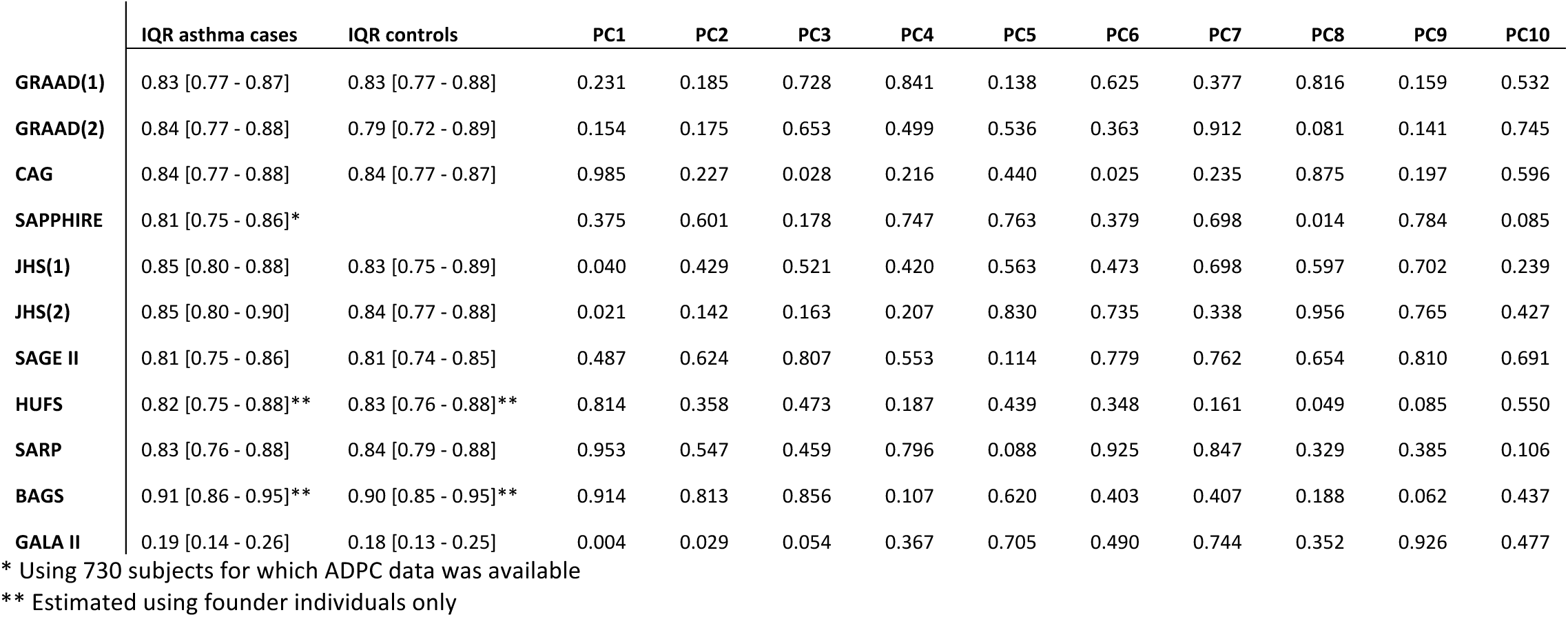
Summary of ancestry distribution in the CAAPA datasets. The median and interquartile range (IQR) of the proportion of African ancestry, estimated using the software program ADMIXTURE, are listed by dataset and asthma case-control status. The p-values of univariate association tests between asthma case-control status and each of the first 10 principal components (PCs) are also listed by study.

**Supplementary Table S13:**
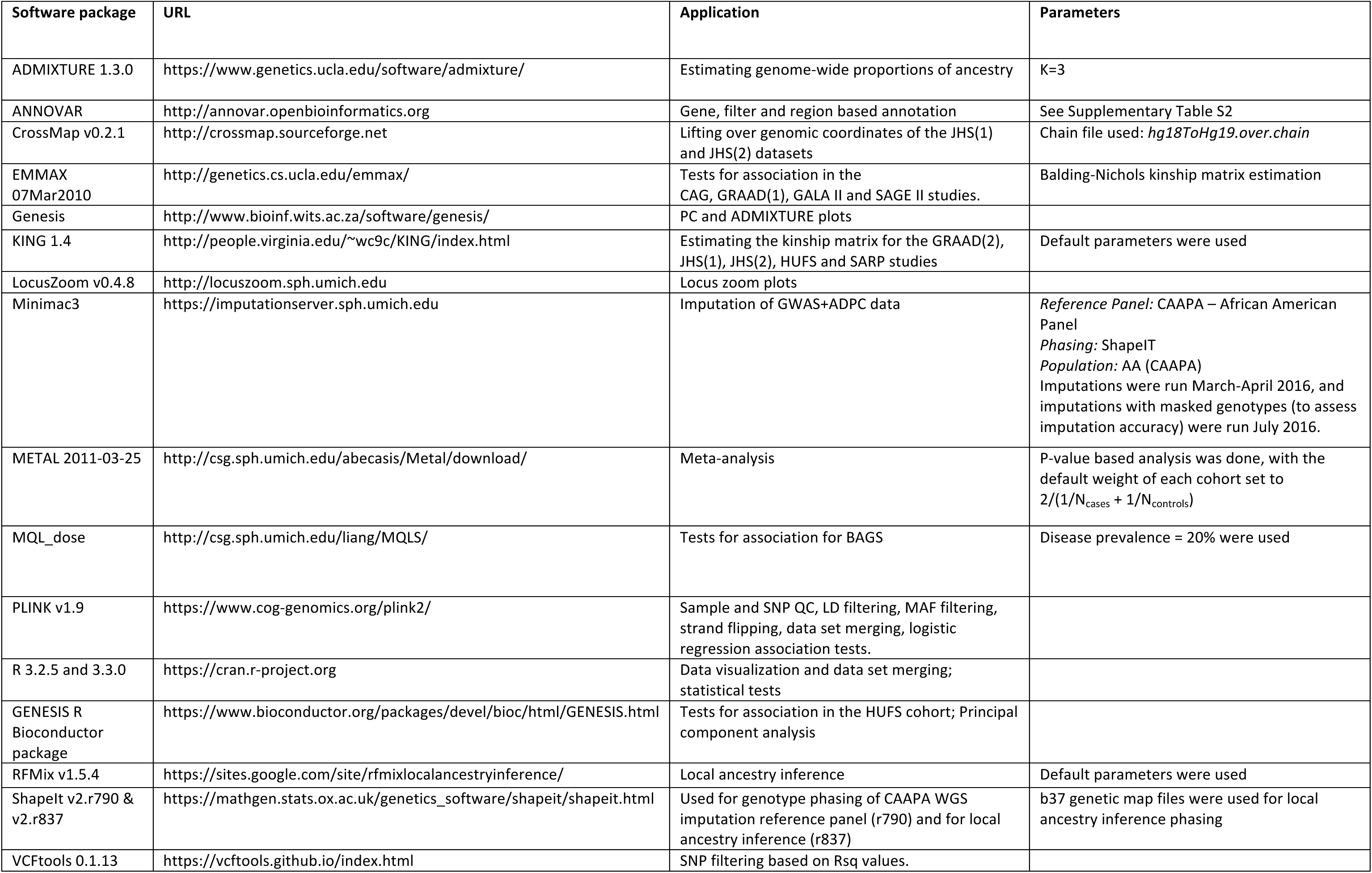
Software used for data analysis. With the exception of software packages used for calling genotype data, this table summarizes the software packages used for processing and analyzing data.

